# Inferring punctuated evolution in the learned songs of African sunbirds

**DOI:** 10.1101/828459

**Authors:** Jay P. McEntee, Gleb Zhelezov, Chacha Werema, Nadje Najar, Joshua V. Peñalba, Elia Mulungu, Maneno Mbilinyi, Sylvester Karimi, Lyubov Chumakova, J. Gordon Burleigh, Rauri C.K. Bowie

## Abstract

Signals used in animal communication, especially those that are learned, are thought to be prone to rapid and/or regular evolution. It has been hypothesized that the evolution of song learning in birds has resulted in elevated diversification rates, as learned song may be subject to especially rapid evolution, and song is involved in mate choice. However, we know little about the evolutionary modes of learned song divergence over timescales relevant to speciation. Here we provide evidence that aspects of the territorial songs of Eastern Afromontane sky island sunbirds *Cinnyris* evolve in a punctuated fashion, with periods of stasis, on the order of hundreds of thousands of years or more, broken up by strong evolutionary pulses. Stasis in learned songs is inconsistent with learned traits being subject to constant or frequent change, as would be expected if selection does not constrain song phenotypes, or if novel phenotypes are frequently advantageous. Learned song may instead follow a process resembling peak shifts on adaptive landscapes. While much research has focused on the potential for rapid evolution in bird song, our results suggest that selection can tightly constrain the evolution of learned songs over fairly long timescales. More broadly, these results demonstrate that some aspects of highly variable, plastic traits can exhibit punctuated evolution, with stasis over fairly long time periods.

## Introduction

Signal evolution has long been thought to be important to the process of animal speciation, in part because many closely related species differ strongly in signals while differing little in other traits (1, 2). In particular, the evolution of signals involved in mate choice has been thought to be critical to the evolution of pre-mating reproductive isolation, such that correlated evolution of signals and mating preferences could lead in and of itself to speciation (3, 4). However, there remain many questions about how signal divergence proceeds over time, which mechanisms are responsible, and how it contributes to speciation and diversification processes.

Some signals that may be important to speciation are highly plastic, including those that are impacted by learning processes (5–7). While divergence in less plastic traits generally requires genetic divergence, the same is not true for learned signals (even if they have components with genetic predispositions (8, 9)). Indeed, novel learned signals can arise without genetic mutation, and spread quickly throughout populations (10). Thus, learned signals are potentially subject to different evolutionary pressures (1, 6), and may exhibit different evolutionary trajectories (11) than signals that are not learned. West-Eberhard (1) suggested that taxa with learned signals may be especially subject to regular and rapid evolutionary change because cultural novelties appear frequently, and because novelty itself may increase the success of new signals.

Modeling has supported the hypothesis that learning can result in increased rates of trait evolution (12). However, selection may not be as effective in driving the evolution of learned signals as compared to more innate signals (13), as learned signals may generally have low heritability (14). In some cases, such a depressed response to selection should lead to slower trait evolution. More generally, modeling of plastic traits has shown that rates of evolution may be fastest at low (15), high (15, 16), or even intermediate (17) levels of plasticity, depending on specific conditions. Thus it remains unclear how quickly learned traits should be expected to evolve relative to traits with less plasticity, and whether learned traits should exhibit similar evolutionary modes (i.e. gradual versus punctuated evolution) to less plastic, non-learned phenotypic traits (11, 13).

The songs of oscine songbirds present intriguing cases for the study of learned signal evolution. In the oscine songbirds, most species learn to perform aspects of songs by imitating conspecifics (18). The oscine learning process is directed by innate predispositions that result in selective learning - that is, species only learn or reproduce vocalizations with certain characteristics (19–21). Song in oscine songbirds typically has two functions - territorial competition and mate attraction (18). While some birds’ vocal repertoires are extensive, a single song component of the vocal repertoire is often used for both of these functions. The mate attraction function of oscine bird song has been especially important to arguments about its relevance to speciation. However, it remains unclear how much of learned song evolution in this group is driven by its role in mate choice, which is central to most hypotheses regarding the role of bird song in speciation (22), compared to territorial competition (23) or other functions.

If song evolution is critical to speciation processes in birds, the evolution of song learning from non-learning ancestors may impact diversification rates in birds. The evolution of song learning, relative to the absence of learning, may decrease the waiting time to speciation (22) and increase diversification rates (24). Alternatively, however, the evolution of song learning may slow the speciation process because learning can result in heterospecific copying between incipient species (25), facilitating hybridization (26). Additionally, song learning is associated with increased within-species variation, which may translate to slower rates in the evolution of song discrimination in incipient species, as larger evolutionary changes to mean phenotypes result in less discrimination in learners than non-learners (27). Slower evolutionary rates of discrimination could result in slower evolutionary rates of prezygotic reproductive isolation, and thus slower speciation rates. Critically, however, we know very little about the trajectories of learned song evolution to inform how learned song evolution may differ from the evolution of less plastic traits.

Previous reviews (1, 11) have suggested that learned signals should exhibit little conservatism, with isolated populations typically diverging before any genetic differences have accrued. Wilkins et al. (11) suggest that non-learned signals, by contrast, may diverge approximately linearly, for example if signal divergence is a function of mutation-order processes. In this way, learned signal divergence might outpace the divergence of non-learned signals early in the divergence process. The strongest evidence for the trajectories of learned song relevant to speciation likely comes from studies of the greenish warbler *Phylloscopus trochiloides*, which exhibits nearly continuous variation across geographic space, suggestive of gradual evolution (28). Gradual evolution has elsewhere been posited to be important in song divergence in birds, with relevance for speciation (29).

Here we examine the evolutionary mode of learned songs in the eastern double-collared sunbird (EDCS) species complex (30), which inhabits mountains of the Eastern Afromontane. Sunbirds are oscine songbirds, and their territorial songs exhibit all signatures of songs developed through learning, including striking complexity and variation (18). The geographic ranges of these species are archipelago-like (Figure 1), with populations occupying discrete, island-like patches of suitable montane forest and forest edge habitats. There is a broad spectrum of molecular divergence, from minimal divergence in some neighboring populations to deep divergences among major lineages.

**Figure 1.**
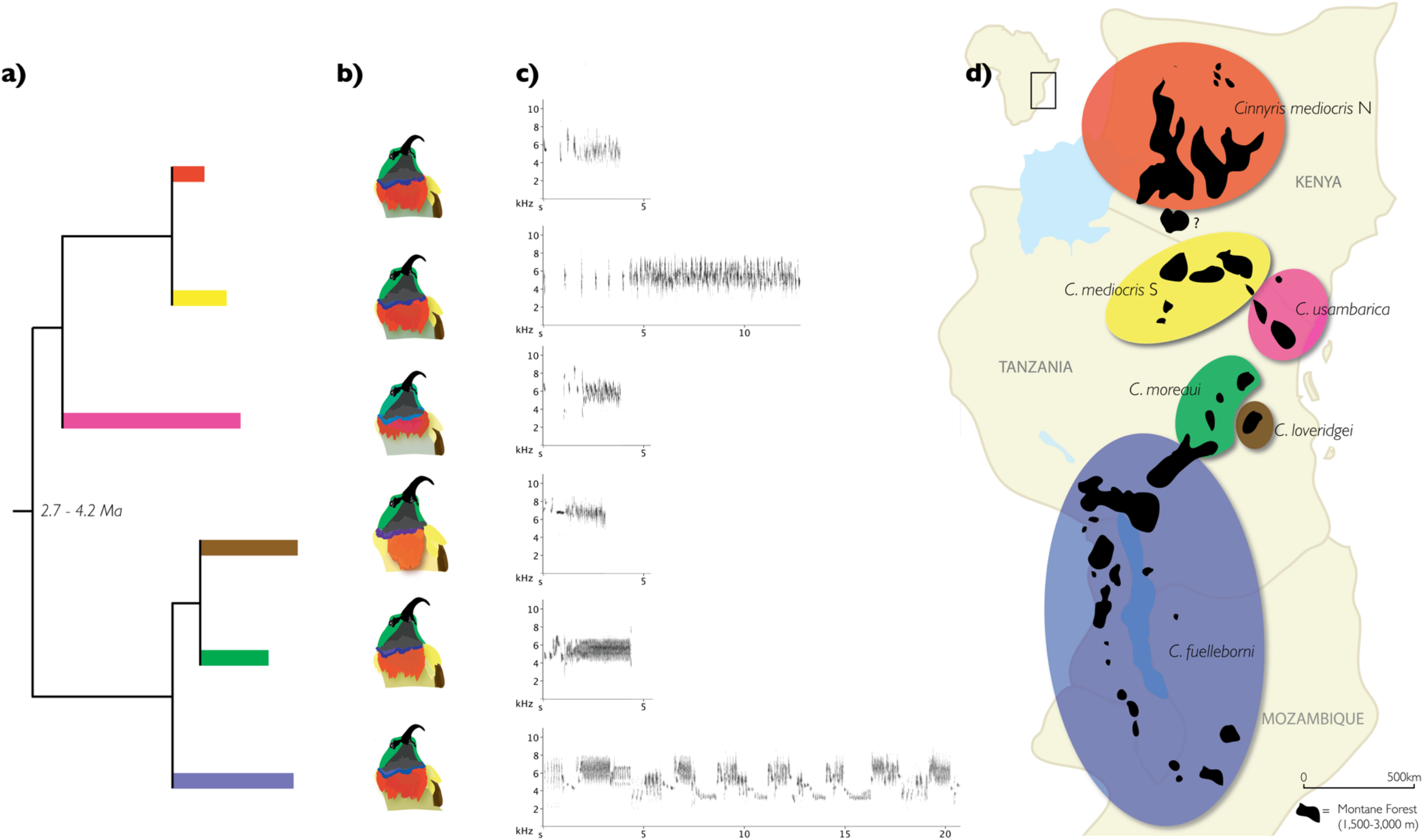
An overview of the Eastern Double-collared Sunbird species complex (EDCS). a) A phylogenetic tree trimmed to include named species and one within-species division that corresponds with a major song divergence. Estimated age of the most recent common ancestor is shown at the node. b) Depictions of typical adult male plumage for the six lineages represented. c) Sonograms showing representative songs for the six lineages shown. d) Ranges of the six lineages in eastern Africa across Kenya, Tanzania, Malawi, and Mozambique.

This species complex allows us to investigate the temporal trajectories of song divergence. Here we first analyze song using multivariate approaches. Then we fit univariate evolutionary models to song traits on phylogenies to examine the tempo and mode of learned song evolution. We were motivated especially by the question of whether learned songs evolve gradually, as is suspected in the Greenish warbler and hypothesized generally for learned bird songs, or via punctuated evolution. This question is rarely posed of signaling phenotypes (31).

## Results

Clustering analyses of 141 multivariate song phenotypes from across the EDCS species complex found support for six distinct phenotypic clusters in multivariate space (the preferred model by BIC had 6 components). Visualization by bivariate plotting demonstrated that the set of phenotypic clusters is separated across different song dimensions - i.e. song divergence across the species complex is multifarious. Some cluster pairs are predominantly separated by frequency, others by fine temporal structure, and others by song duration (Figure A1).

Multi-locus phylogenetic analyses based on the mtDNA gene ND2 and five nuclear intron sequences revealed molecular lineages that correspond with song phenotype clusters (Figure 1, Figures A2 - A3). We recovered five major molecular lineages across the species complex that were similar to those found in previous phylogenetic analyses using only mtDNA sequences (30), and correspond with the taxonomy proposed in Bowie et al. 2004 (which recommended elevating *Cinnyris fuelleborni* and *C. usambarica* to species). Additionally, we recovered distinct clades within three species: *C. mediocris*, *C. fuelleborni*, and *C. moreau*i. In *C. mediocris*, our samples from the Mbulu highlands in northern Tanzania formed a clade, while those from Kenyan populations formed a clade sister to it. *C. fuelleborni* also comprised two clades, with individuals from the Njesi Plateau in northern Mozambique sister to all other *C. fuelleborni*. In *C. moreaui*, samples from the Nguru Mountains formed a clade nested within a phylogenetic grade representing samples from all other localities for this taxon (Figure A3). Phylogenetic analysis using BEAST recovered the same topology for species relationships for the five named species as our ML analysis, and estimated a divergence time of 3.4 My (HPD Interval: 2.67 - 4.18 My) for the most recent common ancestor of the EDCS species complex (Figure A4). Our population trees, which we used to fit evolutionary models for song phenotypes (e.g. Figure 2), recovered the same topology among named taxa as the ML and Bayesian analyses.

**Figure 2.**
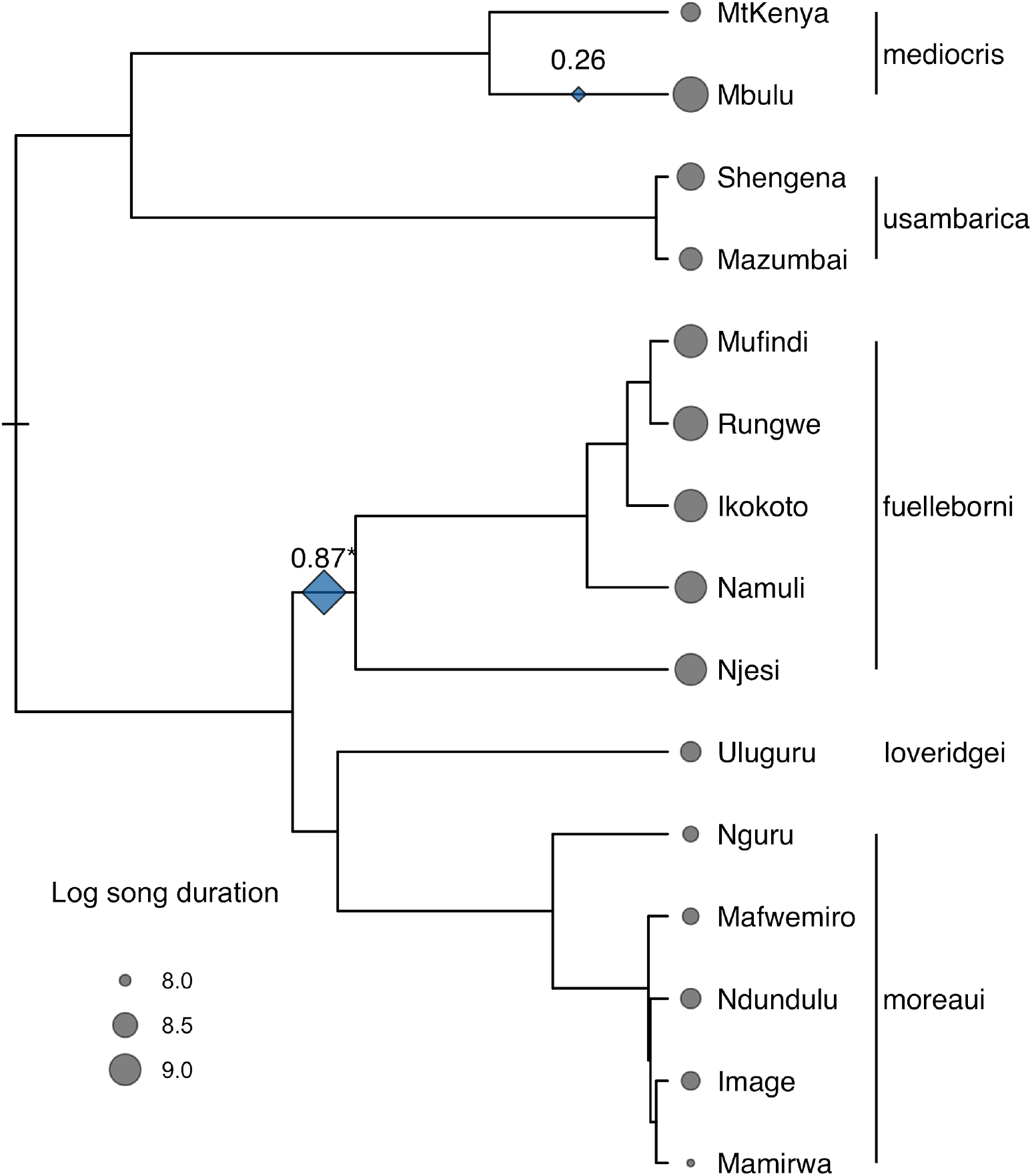
Pulse localization for the evolution of (log) song duration. Blue diamonds are found on branches where pulse localizations had support values > 0.2, with diamond size reflecting support, and support value shown above. The asterisk signifies a pulse that occurs in the pulse configuration with the minimum AICc value. Sizes of gray circles correspond to mean phenotype values at tips, which each represent a geographically discrete sky island population. Species epithets are indicated at far right.

Classification of individuals by song phenotype corresponded closely to the assignment of their populations to major lineages in the molecular phylogenetic tree (with the Mbulu population of *C. mediocris* included as a separate lineage). In unsupervised classification, 93.6% of 141 individuals were assigned correctly to the corresponding molecular lineage (for populations where there were no molecular samples, molecular lineage was presumed based on geography and/or plumage).

### Inference of tempo and mode of learned song evolution

Using novel model-fitting approaches and a novel model implementation for punctuated evolution on phylogenetic trees (see Supplementary Information), we compared support for Brownian motion versus punctuated evolution across different song traits. To compare support, we calculated AICc values for each model. However, we found that standard model comparison in this case was biased to favor punctuated evolution (Supplementary Information), such that we could not report uncorrected ΔAICc weights to represent relative model support. Thus, we performed stochastic simulations under a range of parameters close to the maximum likelihood parameters (within 5%) estimated under both models, then calculated AICc differences for these simulations to calibrate relative support for each model, by each trait. Our approach may be understood as an approximate correction that takes into account phylogenetic correlations present in the data, as is necessary for BIC (32). We found variation across song traits in the relative support for Brownian motion and punctuated evolution. We found strong support for punctuated evolution in four of the fourteen song traits. There was moderate support for punctuated evolution in three more traits. For the remainder, neither model was strongly favored over the other. Sensitivity analyses examining the fit of trait evolution models on bootstrap trees showed that our results were broadly robust to phylogenetic uncertainty (Figure S4).

Our approach allowed us to compare support for the number and positions of evolutionary pulses on our phylogenetic tree. Pulses were allowed to occur on any branch. In our modeling approach, each pulse was considered a parameter, such that more parsimonious models had fewer pulses. We present the results for our pulse localization approach for all song traits where pulsed evolution had strong or moderate support (Figure 2, Figures A4-A9, Table S2) to make evident the degree to which pulses were co-localized across song traits. The pulse configurations with minimum AICc for the seven traits where punctuated evolution was supported had a mean of 1.14±0.35 SD pulse positions (Figure S8). Support for the punctuated evolution model across these traits, coupled with the small number of pulses supported for them, indicated that evolutionary change is minimal for these traits for long stretches of time, corresponding to millions of years, on the phylogenetic tree.

## Discussion

### Learned song evolution as peak shifts on adaptive landscapes

Here we have presented evidence that punctuated evolution explains the evolution of multiple aspects of territorial song better than gradual evolution (Brownian motion), across the EDCS species complex. Our results indicate that the evolutionary mode of multiple aspects of territorial songs includes large jumps in mean trait values, and extended periods of highly bounded evolution, or stasis, in the same aspects. While rapid evolution of animal signals, as in abrupt pulses here, has often been discussed in the literature on signal evolution, and has frequently been invoked as a route to pre-zygotic reproductive isolation in speciation, extended periods of stasis in signals has received comparatively little attention (31). Characterizing the evolutionary mode of territorial song as we have done here sheds light on the form of signal evolution, how it may be involved in speciation processes, and what may or may not cause abrupt evolutionary changes.

The learning process in the development of song in oscine songbirds, like the focal sunbirds here, is a form of phenotypic plasticity (6). As such, our finding that song evolution proceeds as punctuated evolution can be viewed as an example in a learned trait as well as, more generally, in a highly plastic phenotype. As in previous studies using phylogenetic comparative methods (33, 34), we interpret punctuated evolution by visualizing peak shifting on an adaptive landscape. Authors have suggested that phenotypic plasticity itself could assist peak shifting by allowing phenotypes to initially approach alternate peaks on an adaptive landscape without having to wait for novel genetic variation, especially in the case of behavior (35). However, high levels of plasticity may enable so much variation that phenotypes may take on extremely different characteristics without underlying genetic change, such that adaptation to a novel peak does not occur (i.e. plasticity itself is the adaptation). In the case of song, it would seem unlikely that there is a fixed adaptive landscape (16), in which peaks maintain stable shapes, and occupy the same positions through time. Instead, because the efficacy of signals can change depending on environmental variation (e.g. habitat structure (36); population density (37)) or with the evolution of receiver responses (31)), adaptive peaks for learned song would appear likely to change shape, move, appear, and/or disappear, over time and across space. Kirkpatrick (38) and Whitlock (16) showed that even small changes in the slopes and heights of adaptive peaks alone could trigger peak shifts, suggesting they may occur regularly, especially for plastic traits (35). Thus, there are two main theoretical obstacles for highly variable learned song to exhibit peak shifting dynamics over longer timescales. First, song may be so plastic that it can travel about an adaptive landscape without any underlying genetic change (35, 39), in which case it would likely be prone to extremely high lability. Secondly, the adaptive landscape for song may change rapidly through time, and vary across environments, such that adaptive peaks are unlikely to remain in the same shape and position over evolutionary timescales (16, 31). Thus, there was little reason to expect learned songs to be restricted to peaks, because of their high variability, and little reason to expect adaptive peaks to be stable in position and shape over time such that they could be observed.

Our finding that punctuated evolution better characterizes the evolution of some song traits than gradual evolution indicates that evolution can be tightly bounded, approaching stasis, over long periods of time (> 10^6^ years). These results suggest that adaptive peaks for song are stable over time. The stability of adaptive peaks for learned songs suggests that the songs’ receivers mediate stabilizing selection on song traits. There are two sets of receivers, males and females, that are likely to exert stabilizing selection forces in sunbirds. If narrow female preferences alone were responsible for stasis, we would expect strong behavioral reproductive isolation where two species with highly divergent song come into contact. However, *C. moreaui* and *C. fuelleborni*, which have extremely different songs across many song aspects, hybridize where they come into contact (40), indicating that female preferences are unlikely to be narrow. Thus, male receivers are likely to play an important role in the evolutionary stasis of learned song traits, by exerting quadratic selection itself, by exerting directional selection opposite selection from females, or perhaps by exerting selection opposite natural selection. An alternate hypothesis for stasis in some traits is that evolutionary constraints result from limited genetic variance (41–44). However, limited genetic variance should not strongly constrain evolution over longer timescales, as examined here, because novel genetic variation will arise over these timescales.

If near-stasis occurs over long periods of time in some bird song traits, what explains evolutionary divergence when it occurs? One prominent hypothesis explaining the evolution of bird song is that song evolves as a byproduct of morphological evolution. Two aspects of morphology have been consistently highlighted with respect to such byproduct evolution: beak morphology and body size. Morphological evolution of the beak may be important as the beak is a part of the vocal apparatus itself (45). Body size evolution may be important because of allometric changes in pieces of the vocal apparatus, which could alter song frequency (46). When song evolves by punctuated evolution, the morphological byproduct hypothesis would predict that evolutionary pulses are consequences of morphological evolution (which in itself might be punctuated). In the EDCS, there is limited morphological evolution, with subtle changes in morphology across the complex, and substantial overlap in morphological characteristics that differ on average between species (30). Thus, there is overall little reason to suspect that song evolution is tightly connected to morphological evolution. However, Loveridge’s Sunbird *C. loveridgei* is unambiguously the largest member of the species complex, and has the longest bill. Thus this species represents a test case for the predictions of the morphological by-product hypothesis for acoustic signal evolution, in which we would predict that Loveridge’s Sunbird should have the lowest frequency songs within the species complex. We find the opposite of this prediction. Loveridge’s Sunbirds sing songs with the highest peak frequencies of all the members of the species complex, and our analyses evince a pulse of peak frequency evolution unique to Loveridge’s Sunbird. Thus, within the EDCS complex, we see punctuated evolution that is not associated with substantial morphological evolution generally, and in the instance where morphological divergence is most pronounced, song evolution exhibits pulsed change opposite the direction predicted under the by-product hypothesis. Thus the overall picture from this species complex is that song evolution is not contingent on morphological evolution, extensive song diversity is generated without great morphological change, and the impacts of morphological evolution on song evolution are dwarfed by other processes.

Range expansion provides another possibility as a cause for pulses in learned songs. Studies on North American juncos (47, 48) have suggested that pulses of phenotypic divergence (in that case, plumage) might take place in association with instances of rapid range expansion. During range expansion, serial founder effects can induce the fixation of rare genetic variants, providing a mechanism for rapid genetic change. Moreover, selective forces on signals may be distinct at the leading front of range expansions. For example, population densities at the leading edges of range expansions may be low, which could advantage signals that broadcast across further distances. In the future, genome-wide molecular studies could be used to reconstruct range expansions to examine correspondence in phenotypic change with range expansion in the EDCS. Studies of the transmission properties of the different song phenotypes across the species complex are also desirable, as they could inform hypotheses about song evolution based on population densities.

As a learned trait, bird song may be prone to evolution via genetic assimilation of phenotypic novelties without an initial genetic basis (5, 6). Traits showing pulsed evolution - frequency, fine-scale temporal structure, duration - are traits that have been shown to have underlying genetic variation in other songbirds (8, 9, 39). Based on this evidence, we suspect that these traits have underlying genetic predispositions in the focal species complex. This hypothesis is supported by the absence of a cultural bleed of phenotypes across the *C. moreaui* - *C. fuelleborni* contact zone where the two species interact (40). Genetic assimilation remains a plausible path for evolutionary divergence (i.e. “genes as followers” (5)), as underlying genetic differences are the end stage of genetic assimilation. However, peak shifting mechanisms involving substantial genetic change concomitant with phenotypic change are also plausible in explaining song evolution here, as we do not find evidence for substantial song divergence in the absence of genetic differentiation.

### Relevance of punctuated evolution of learned song for speciation

The EDCS species complex bears hallmarks of speciation by sexual (2), or social (1) selection: species are strongly divergent for a signal used in social competition, and do not differ strongly in ecological respects (40). Panhuis et al. (2) suggested that an additional signature of speciation by sexual selection is the evolution of variation in sexually selected traits among populations within species, with this variation generating partial premating isolation. Our sampling of isolated sky island populations, especially within *C. moreaui* and *C. fuelleborni*, allows us to characterize within-species variation in territorial song. Across most song traits, variation across populations within species is minimal, including for many traits with strong differences across species, e.g. CV peak frequency (Figure A5) and median pause duration (Figure A7). As such, between-species divergence cannot be extrapolated from within-species variation (49). There are discontinuities in evolutionary processes in learned song that give rise to the diversity of songs across the species complex. These discontinuities in evolutionary process appear responsible for species differences.

## Conclusions

The effects of learning on evolutionary processes are poorly known. Previous work has suggested that stabilizing selection on learned traits should be inadequate to prevent the divergence of genetic predispositions by drift, ultimately facilitating more rapid divergence in those genes underlying traits (22). Our study shows that multiple song traits can exhibit stasis for prolonged periods, likely lasting hundreds of thousands of years or more. These results suggest that learned song in the focal taxa is subject to a combination of sufficiently strong stabilizing selection and sufficient exposure of the underlying genetic variation to prevent incremental change for long periods of time. An alternative, that there is insufficient genetic variation underlying these traits, is potentially plausible, but appears less likely given the evidence that genetic variation for learned song traits is present in captive populations, and the long span of evolutionary time during which such variation could be generated.

## Materials and methods

### Song analysis

We made sound recordings of EDCS from 2007-2011 in Kenya, Tanzania, and Mozambique, using solid-state digital recorders (Marantz PMD models 660, 661, and 670) and shotgun microphones (Sennheiser ME-67). A small number of recordings were made using a parabolic dish with an omnidirectional microphone (Sennheiser ME-62). We complemented our field recordings with additional recordings from the Macaulay Library (http://macaulaylibrary.org) and the British Library of Natural Sounds (https://www.bl.uk/collection-guides/wildlife-and-environmental-sounds). The vocal repertoires of the focal taxa are complex, including a wide array of different signal types. Here we measure the acoustic properties of male territorial songs delivered in bout form, in which consecutive songs are typically separated by a short duration (<15 s) of silence, or a series of short calls and pauses (50). Sunbirds sing these songs from a perch in the vegetation, ranging in height from 2 to 30m. These songs function in male-male territorial interactions (51). Further, as in other passerine birds (18), these songs likely serve to attract mates. Singing can coincide with, or immediately precede, female wing-fluttering displays directed at singing males, which has been observed in *C. loveridgei* and *C. fuelleborni* (JPM pers. obs.).

Before analyses, recordings were standardized for frequency sampling at 44.1 kHz, and bandpass filtered at 2 to 10 kHz. More strict filtering, at 2.5 to 9kHz, was then employed for recordings of *C. mediocris* and *C. usambarica* to allow fine-scale structural analysis of sonograms, as our recordings of their songs generally had lower signal:noise ratios, and the lowest frequencies in their songs are >2.5 kHz. Similarly strict filtering could not be applied to *C. fuelleborni* or *C. moreaui* songs because their songs sometimes include peak frequencies below 2.5kHz. Spot filtering was used to remove acoustic signals not emitted by the focal bird. We selected high-quality field recordings for analyses after sonogram visualization in Raven Pro 1.3 (52). JPM performed all sonogram analysis procedures in the program Luscinia (53). Sonograms were produced in Luscinia with the following settings: maximum frequency: 10 kHz; frame length 5 ms; time step: 1 ms; spectrograph points: 221; spectrograph overlap: 80%; echo removal: 100%; echo range: 100; windowing function: Hann; and high pass threshold: 2 kHz. Signals within sonograms were detected using Luscinia’s automated signal detection. Results of automatic signal detection procedures were checked by eye and ear, with recordings slowed for playback to ⅛ speed. Automated signal detection errors were corrected using the *brush* tool. Measurements were made for each sonogram trace (hereafter ‘elements’), separated by pauses from other elements.

From the set of measurements of each element, we calculated summary statistics at the song level. For each individual sunbird, we then calculated the mean values of a set of summary statistics across songs. We calculated the following summary statistics for each song, based on values for each element: median pause duration between elements (ms), coefficient of variation (cv) of pause duration (ms), median peak frequency (Hz), cv peak frequency, maximum peak frequency (Hz), minimum peak frequency (Hz), range peak frequency (difference between maximum and minimum peak frequencies), number of elements, median frequency bandwidth (Hz), cv frequency bandwidth (Hz), median frequency change (Hz), cv per-element frequency change (Hz), song duration (ms), and median element duration (ms). Peak frequency is defined as the frequency window with the highest amplitude for a given portion of the sonogram. We took the natural log of the number of elements, median frequency change, median frequency bandwidth, and song duration to improve downstream analyses with respect to assumptions of normality. To generate estimates of song phenotypes at the level of the individual bird, we took the arithmetic mean of the values for each variable across songs. These procedures resulted in a data set comprising song phenotype estimates for 142 individuals from measurements of 419 songs. A mean of 2.95±1.01SD songs were measured per individual.

We used Gaussian finite mixture modeling (GFMM) to perform cluster analyses on the 14 song traits measured for each individual. GFMM was performed using the package Mclust (54) in R 3.5.2 (55). We built models with the number of mixture components varying from 1 to 9, and interpreted each of these components as a cluster in multivariate phenotypic space (56). For each number of specified components, we built six different types of models representing different parameterizations of the covariance matrix. These parameterizations allow for flexibility in the volume and shape of the mixture components. We examined relative support for the 54 total models using the Bayesian Information Criterion (BIC).

### Molecular phylogenetics

We performed phylogenetic analyses using DNA sequence data for samples collected from the field (see the Appendix for details on sampling for molecular analyses and for further detail on phylogenetic methods, see SI Appendix 2 for specimen details). First, to investigate whether song phenotypes generally correspond to phylogenetic lineages across the species complex, we built a multi-locus phylogenetic tree from a concatenated alignment of DNA sequences for three mtDNA genes (ND2, ND3, and ATP6) and six nuclear autosomal introns (MB, CHDZ, 11836, 18142, TGFb2, and MUSK), for 256 in-group individuals and 12 outgroup species. The alignment had 5,313 base pairs. We estimated our phylogenetic tree using a maximum likelihood approach (57), and assessed support for nodes by bootstrapping. Secondly, we built population-level trees to investigate the history of song and beak length divergence over the focal species complex. We constructed these trees by calculating mean population distances for 16 and 15 populations, for analyses of song and beak phenotypes respectively. For these trees, we used sequences of the mtDNA genes ND2 (440 bp) and ND3 (362 bp), for 134 and 128 individuals for song and beak analyses, respectively. Topological relationships between named species were the same in the mtDNA population trees as in the multi-locus phylogenetic tree with species as tips.

Third, we sought to estimate the age of the most recent common ancestor of the species complex. We performed a species-level phylogenetic analysis using a Bayesian coalescent-based method (BEAST, (58, 59)) with DNA sequence data from two mtDNA genes (ND2 and ATP6, coded as a single locus) and four nuclear DNA sequences (ATP6, TGFb2, MB, and CHDZ). This analysis included 16 species as tips, including the five named species in the focal species complex (*C. mediocris*, *C. usambarica*, *C. loveridgei*, *C. moreaui*, and *C. fuelleborni*), eight other sunbird species, and three species of flowerpecker (Dicaeidae). We dated the most recent common ancestor (MRCA) of the focal species complex by implementing a normal prior distribution (mean = 18 My, SD = 2) on the node age of the MRCA of sunbirds and flowerpeckers, based on a recent dating analysis of a family-level phylogenetic tree of the Passeriformes (60). We used a GTR-gamma substitution model and a relaxed lognormal molecular clock, with substitution rate prior distributions based on divergence rate estimates for the Hawaiian honeycreeper radiation (an oscine passerine radiation, like sunbirds) (61).

### Phylogenetic comparative method approach

To investigate the tempo and mode of song divergence, we compared phylogenetic trait evolution models fit to population-level data. We fit models to each song trait individually. We built and fit phylogenetic trait evolution models representing 1) strongly bounded evolution (stasis or near-stasis) punctuated by pulses and 2) gradual evolution (Brownian motion), by maximum likelihood. These models are described in the Supplementary Information. Models including pulses were fit under the condition that there were a maximum of four pulses across the population tree. We compared support for fitted models using AICc values (62). To characterize the uncertainty in model selection due to phylogenetic uncertainty, we built ten bootstrap population trees, and fit both trait evolution models on each of the bootstrap phylogenies for each song trait.

For the traits where punctuated evolution models were a better fit than Brownian motion, we estimated the location of pulses on the population tree. Our method for fitting pulsed evolutionary models involves fitting the maximum likelihood parameters for all potential pulse configurations on the tree, given a maximum of four pulses (corresponding to ~4.6 × 10^4^ pulse configurations). To quantify the strength of the evidence for an evolutionary pulse on a given branch of the phylogenetic tree, we calculated the sum of the AICc weights of those pulse configurations that include a pulse on the given branch, and divided this by the sum of the AICc weights of all computed pulse configurations (63). Our approach treats each *n*-pulse configuration as an independent mode We present maximum likelihood parameters for the maximum likelihood configuration (Tables S1 and S2).

## Supporting information

sampling for molecular analyses

## Acknowledgements

We thank Jessica Hughes, Violet Kimzey, Somin Lim, Kyle Marsh, Dalila Sequeira, Emilia Wakamatsu, Cynthia Wang, and Addien Wray for research assistance. Kathleen Rudolph assisted by designing and illustrating Figure 1. Ruth King contributed advice on statistical issues. David Moyer, Norbert Cordeiro, Liz Baker, and Neil Baker provided advice and material support for field work. Caffe Luce in Tucson, Arizona, provided work space. Funding was provided by the National Geographic Society; Waitt Foundation; Mohamed bin Zayed Species Conservation Fund; a National Science Foundation (NSF) Doctoral Dissertation Improvement Grant (Award #1011702); a Beim Summer Research Award (UC Berkeley Dept. of Integrative Biology), Lousie Kellogg, Albert Preston Hendrickson, and Charles Koford grants from the Museum of Vertebrate Zoology; a UC Berkeley Center for African Studies Andrew and Mary Thompson Rocca Scholarship; American Ornithologists’ Union (now American Ornithological Society); the Explorers Club; and NSF grant DBI-1458034. GZ and LC were supported by the Leverhulme trust grant RPG-2017-249. GZ was supported by NSF grant DMS-1056471. Research permissions for field work were furnished by: TAWIRI, COSTECH, and the Ministry of Natural Resources, Forestry and Beekeeping Division, Tanzania; National Museum Kenya, National Council for Science and Technology, and Kenya Wildlife Service, Kenya; and the Universidade Eduardo Mondlane Natural History Museum of Maputo and Ministry of Agriculture, Mozambique.

## References

1. M. J. West-Eberhard, Sexual Selection, Social Competition, and Speciation. Q. Rev. Biol. 58, 155–183 (1983).

2. T. M. Panhuis, R. Butlin, M. Zuk, T. Tregenza, Sexual selection and speciation. Trends Ecol. Evol. 16, 364–371 (2001).

3. R. Lande, Models of speciation by sexual selection on polygenic traits. Proc. Natl. Acad. Sci. U.S.A. 78, 3721–3725 (1981).

4. M. Kirkpatrick, Sexual selection and the evolution of female choice. Evolution 36, 1–12 (1982).

5. M. J. West-Eberhard, Developmental plasticity and the origin of species differences. Proc. Natl. Acad. Sci. U.S.A. 102, 6543–6549 (2005).

6. M. J. West-Eberhard, Developmental plasticity and evolution (Oxford University Press, 2003).

7. M. N. Verzijden, et al., The impact of learning on sexual selection and speciation. Trends Ecol. Evol. 27, 511–519 (2012).

8. P. C. Mundinger, D. C. Lahti, Quantitative integration of genetic factors in the learning and production of canary song. Proceedings of the Royal Society B - Biological Sciences 281, 20132631 (2014).

9. D. G. Mets, M. S. Brainard, Genetic variation interacts with experience to determine interindividual differences in learned song. Proc. Natl. Acad. Sci. U.S.A. 115, 421–426 (2018).

10. R. B. Payne, L. L. Payne, S. M. Doehlert, Biological and cultural success of song memes in Indigo Buntings. Ecology 69, 104–117 (1988).

11. M. R. Wilkins, N. Seddon, R. J. Safran, Evolutionary divergence in acoustic signals: causes and consequences. Trends Ecol. Evol. 28, 156–166 (2013).

12. G. E. Hinton, S. J. Nowlan, “How learning can guide evolution” in Adaptive Individuals In Evolving Populations: Models And Algorithms, (Addison-Wesley Reading, MA, 1996), pp. 447–454.

13. I. Paenke, B. Sendhoff, T. J. Kawecki, Influence of plasticity and learning on evolution under directional selection. Am. Nat. 170, E47–58 (2007).

14. T. D. Johnston, “Selective costs and benefits in the evolution of learning” in Advances in the Study of Behavior, J. S. Rosenblatt, R. A. Hinde, C. Beer, M.-C. Busnel, Eds. (Academic Press, 1982), pp. 65–106.

15. E. Borenstein, I. Meilijson, E. Ruppin, The effect of phenotypic plasticity on evolution in multipeaked fitness landscapes. J. Evol. Biol. 19, 1555–1570 (2006).

16. M. C. Whitlock, Founder effects and peak shifts without genetic drift: Adaptive peak shifts occur easily when environments fluctuate slightly. Evolution 51, 1044–1048 (1997).

17. R. French, A. Messinger, Genes, phenes and the Baldwin effect: Learning and evolution in a simulated population in Artificial Life IV, (MIT Press, 1994), pp. 277–282.

18. C. K. Catchpole, P. J. B. Slater, Bird Song (Cambridge University Press, 2008).

19. J. A. Soha, P. Marler, A species-specific acoustic cue for selective song learning in the white-crowned sparrow. Anim. Behav. 60, 297–306 (2000).

20. P. Marler, S. Peters, Selective vocal learning in a sparrow. Science 198, 519–521 (1977).

21. P. Marler, S. Peters, The role of song phonology and syntax in vocal learning preferences in the song sparrow, Melospiza melodia. Ethology 77, 125–149 (1988).

22. R. F. Lachlan, M. R. Servedio, Song learning accelerates allopatric speciation. Evolution 58, 2049–2063 (2004).

23. R. M. Tinghitella, et al., On the role of male competition in speciation: a review and research agenda. Behav. Ecol. 29, 783–797 (2017).

24. N. A. Mason, et al., Song evolution, speciation, and vocal learning in passerine birds. Evolution 71, 786–796 (2017).

25. J. Vokurková, et al., The causes and evolutionary consequences of mixed singing in two hybridizing songbird species *(Luscinia* spp.). PLoS One 8, e60172 (2013).

26. A. Qvarnstrom, J. Haavie, S. A. Saether, D. Eriksson, T. Part, Song similarity predicts hybridization in flycatchers. J. Evol. Biol. 19, 1202–1209 (2006).

27. B. G. Freeman, G. A. Montgomery, D. Schluter, Evolution and plasticity: Divergence of song discrimination is faster in birds with innate song than in song learners in Neotropical passerine birds. Evolution 71, 2230–2242 (2017).

28. D. E. Irwin, Song variation in an avian ring species. Evolution 54, 998–1010 (2000).

29. S. V. Edwards, et al., Speciation in birds: Genes, geography, and sexual selection. Proc. Natl. Acad. Sci. U.S.A. 102, 6550–6557 (2005).

30. R. C. K. Bowie, J. Fjeldsa, S. J. Hackett, T. M. Crowe, Systematics and biogeography of double-collared sunbirds from the Eastern Arc Mountains, Tanzania. Auk 121, 660–681 (2004).

31. S. J. Arnold, L. D. Houck, Can the Fisher-Lande process account for Birds of Paradise and other sexual radiations? Am. Nat. 187, 717–735 (2016).

32. M. Khabbazian, R. Kriebel, K. Rohe, C. Ané, Fast and accurate detection of evolutionary shifts in Ornstein–Uhlenbeck models. Methods Ecol. Evol., 811–824 (2017).

33. M. J. Landis, J. G. Schraiber, Pulsed evolution shaped modern vertebrate body sizes. Proc. Natl. Acad. Sci. U.S.A. (2017) https:/doi.org/201710920.

34. M. A. Butler, A. A. King, Phylogenetic comparative analysis: A modeling approach for adaptive evolution. Am. Nat. 164, 683–695 (2004).

35. T. D. Price, A. Qvarnström, D. E. Irwin, The role of phenotypic plasticity in driving genetic evolution. Proc. Biol. Sci. 270, 1433–1440 (2003).

36. H. Slabbekoorn, T. B. Smith, Habitat-dependent song divergence in the little greenbul: An analysis of environmental selection pressures on acoustic signals. Evolution 56, 1849–1858 (2002).

37. E. S. C. Scordato, Male competition drives song divergence along an ecological gradient in an avian ring species. Evolution 72, 2360–2377 (2018).

38. M. Kirkpatrick, Quantum evolution and punctuated equilibria in continuous genetic characters. Am. Nat. (1982).

39. W. Forstmeier, C. Burger, K. Temnow, S. Derégnaucourt, The genetic basis of zebra finch vocalizations. Evolution 63, 2114–2130 (2009).

40. J. P. McEntee, et al., Social selection parapatry in Afrotropical sunbirds. Evolution 70, 1307–1321 (2016).

41. E. B. Kruuk, et al., Antler size in red deer: heritability and selection but no evolution. Evolution 56, 1683–1695 (2002).

42. M. W. Blows, A. A. Hoffmann, A reassessment of genetic limits to evolutionary change. Ecology 86, 1371–1384 (2005).

43. M. W. Blows, S. F. Chenoweth, E. Hine, Orientation of the genetic variance-covariance matrix and the fitness surface for multiple male sexually selected traits. Am. Nat. 163, 329–340 (2004).

44. E. Hine, S. F. Chenoweth, M. W. Blows, Multivariate quantitative genetics and the lek paradox: genetic variance in male sexually selected traits of Drosophila serrata under field conditions. Evolution 58, 2754–2762 (2004).

45. J. Podos, S. K. Huber, B. Taft, Bird song: The interface of evolution and mechanism. Annu. Rev. Ecol. Evol. Syst. 35, 55–87 (2004).

46. M. J. Ryan, E. A. Brenowitz, The role of body size, phylogeny, and ambient noise in the evolution of bird song. Am. Nat. 126, 87–100 (1985).

47. G. Friis, P. Aleixandre, R. Rodríguez-Estrella, A. G. Navarro-Sigüenza, B. Milá, Rapid postglacial diversification and long-term stasis within the songbird genus *Junco:* phylogeographic and phylogenomic evidence. Mol. Ecol. 25, 6175–6195 (2016).

48. B. Mila, J. E. McCormack, G. Castaneda, R. K. Wayne, T. B. Smith, Recent postglacial range expansion drives the rapid diversification of a songbird lineage in the genus Junco. Proceedings of the Royal Society B-Biological Sciences 274, 2653–2660 (2007).

49. P. Campbell, et al., Geographic variation in the songs of Neotropical singing mice: Testing the relative importance of drift and local adaptation. Evolution 64, 1955–1972 (2010).

50. J. P. McEntee, “Social selection, song evolution, and the ecology of parapatry in sunbirds,” University of California, Berkeley, Berkeley, California. (2013).

51. J. P. McEntee, Reciprocal territorial responses of parapatric African sunbirds: Species-level asymmetry and intraspecific geographic variation. Behav. Ecol. 25 (2014).

52. Bioacoustics Research Program, Raven Pro: Interactive Sound Analysis Software (Version 1.3) (2008).

53. R. Lachlan, Luscinia (2007).

54. C. Fraley, A. E. Raftery, T. B. Murphy, “mclust Version 4 for R: Normal Mixture Modeling for Model-Based Clustering, Classification, and Density Estimation” (University of Washington, 2012).

55. R Core Team, R: A language and environment for statistical computing (2012).

56. J.-P. Baudry, A. E. Raftery, G. Celeux, K. Lo, R. Gottardo, Combining mixture components for clustering. J. Comput. Graph. Stat. 19, 332–353 (2010).

57. A. Stamatakis, RAxML version 8: a tool for phylogenetic analysis and post-analysis of large phylogenies. Bioinformatics 30, 1312–1313 (2014).

58. A. J. Drummond, A. Rambaut, BEAST: Bayesian Evolutionary Analysis by Sampling Trees. BMC Evol. Biol. 7, 214 (2007).

59. A. J. Drummond, M. A. Suchard, D. Xie, Bayesian phylogenetics with BEAUti and the BEAST 1.7. Mol. Biol. (2012).

60. C. H. Oliveros, et al., Earth history and the passerine superradiation. Proc. Natl. Acad. Sci. U.S.A. (2019) https:/doi.org/10.1073/pnas.1813206116.

61. H. R. L. Lerner, M. Meyer, H. F. James, M. Hofreiter, R. C. Fleischer, Multilocus resolution of phylogeny and timescale in the extant adaptive radiation of Hawaiian honeycreepers. Curr. Biol. 21, 1838–1844 (2011).

62. K. P. Burnham, D. R. Anderson, Model Selection and Multimodel Inference: A Practical Information-Theoretic Approach (Springer Science & Business Media, 2003).

63. S. T. Buckland, K. P. Burnham, N. H. Augustin, Model selection: an integral part of inference. Biometrics, 603–618 (1997).

