## Appendix 1 for "Inferring punctuated evolution in the learned songs of African sunbirds"

**Appendix 1: Inferring punctuated evolution in the learned songs of African sunbirds**

**Authors:**

Jay P. McEntee[1,2*], Gleb Zhelezov[3], Chacha Werema[4], Nadje Najar[5], Joshua V. Peñalba[6], Elia Mulungu[7], Maneno Mbilinyi[8], Sylvester Karimi[9], Lyubov Chumakva[3], J. Gordon Burleigh[2], & Rauri C.K. Bowie[1]

**Author Affiliations:**

[1]Museum of Vertebrate Zoology & Department of Integrative Biology, University of California, Berkeley, Berkeley, CA 94720 USA

[2]Biology Department, University of Florida, Gainesville, FL 32608 USA

[3]School of Mathematics, University of Edinburgh, Edinburgh EH9 3FD UK

[4]Department of Zoology and Wildlife Conservation, P.O. Box 35064, University of Dar-es-salaam, Tanzania

[5]School of Natural Resources, University of Nebraska, Lincoln. Lincoln, NE 68503 USA

[6]Division of Evolutionary Biology, Ludwig-Maximilians-Universität Munchen, Faculty of Biology, Biocenter, Großhaderner Str. 2, 82152 Planegg-Martinsried, Germany

[7]P.O. Box 934, Iringa, Tanzania

[8]Tanzania Bird Atlas, Iringa, Tanzania

[9]National Museums Kenya, Nairobi, Kenya

### Phylogenetic methods

For phylogenetic analyses, we performed novel DNA sequencing in addition to using sequences previously published. For the maximum likelihood phylogenetic approach, we sought to include sequences from across the Eastern Double-collared Sunbird species complex, as well as for a set of outgroup taxa. For the population tree approach, we sought to maximize sampling for those populations for which we had best sampling of song recordings. For the Bayesian coalescent phylogenetic analyses used to estimate divergence times, we included sequences for a single individual per taxon.

#### *DNA sequencing*

We extracted DNA from tissues and blood samples using Qiagen DNeasy extraction kits (Valencia, CA, USA) and associated standard protocols. Novel DNA sequencing for this study involved PCR amplification for three mtDNA genes (NADH2 ((Sorenson et al. 1999)), NADH3 + flanking tRNAs (Chesser 1999), and ATP6 (Hunt et al. 2001)), four nuclear autosomal introns (MB intron-2 and TGFb2 intron-5 (Kimball et al. 2009); 26S proteasome non-ATPase regulatory subunit 6: here referred to as 11836, and proteasome 26S subunit 14: here referred to as 18142 (Backström et al. 2008)), and two introns from the Z (sex-linked) chromosome (CHDZ intron-15

(Griffiths and Korn 1997) and MUSK intron-3 (Kimball et al. 2009)). We used standard protocols for thermocycling, then purified PCR products with shrimp phosphatase and exonuclease (exoSAPit, Amersham, Foster City, CA, USA). Cycle sequencing was performed in both directions using a BigDye Terminator V3 chemistry (Applied Biosystems, Inc., Foster City, CA, USA) approach to label bases. We performed sequencing on an AB3100 DNA sequencer (Applied Biosystems, Inc., Foster City, CA, USA).

We examined chromatograms for quality in Sequencher 4.7 (Gene Codes Corporation) and Geneious Pro 5.1.6, and checked mtDNA genes for stop codons, insertions, and deletions to prevent the inclusion of pseudogene sequences. We performed alignments using the MAFFT algorithm (Katoh et al. 2005), and then made refinements to alignments by eye. Novel sequences for this study have been deposited in GenBank (accession numbers to be determined).

*Maximum likelihood phylogenetic reconstruction*

To construct a phylogenetic hypothesis for the Eastern Double-collared Sunbird species complex (EDCS), we built an alignment for 174 EDCS individuals and 12 other species (before removing identical sequences, there were 256 in-group individuals and 12 outgroup species). Our alignment for this analysis included sequences from all loci used in this study: three mtDNA genes (ATP6: 675 bp, NADH2: 1,041 bp, and NADH3 + flanking tRNAs: 395 bp), four nuclear autosomal introns (MB: 806 bp, 11836: bp, 18142: bp, and TGFb2: 641 bp), and two Z-chromosome introns (CHDZ: 551 bp and MUSK: 499 bp). Sequence coverage across individuals was highly heterogeneous, as we included all individuals sequenced for any of the genes included. In our maximum likelihood analysis, the proportion of gaps and completely undetermined characters in the alignment was 73.12%.

We used model selection in PartitionFinder 2 to select a partitioning scheme for nucleotide substitution models across sites in our alignment (Lanfear et al. 2012, 2017). We included general-time-reversible (GTR) models, GTR models with gamma-distributed rate variation (GTR+G), and GTR+G models with invariable sites GTR+I+G as candidate nucleotide substitution models across the alignment. We then allowed each model subset (partition) to have distinct base frequencies, rate matrices, and proportions of invariant sites. For mtDNA genes, we allowed each codon position for each gene to have unique model subset parameters. For nuclear introns, we allowed each intron to have unique model subset parameters. We used the “greedy” algorithm in PartitionFinder 2 to search across partition schemes, and we judged support for nucleotide substitution models by comparing AICc values (Burnham and Anderson 2003). The preferred model was a GTR+G model with six partitions.

We performed a joint bootstrap analysis and maximum likelihood tree search using RAxML 8.2.10 (Stamatakis 2014), on the CIPRES Science Gateway (Miller et al. 2010). In these analyses, we specified the clade including the flowerpecker species *Dicaem celebicum*, *Dicaeum agile*, and *Prionochilus maculatus* as the outgroup. We performed 1000 bootstrap replicates, and report bootstrap support values for nodes of interest in Figure S2.

*Population trees*
To generate population trees to be used in evolutionary modeling of traits, we sequenced the mtDNA genes NADH2 and NADH3, and built an alignment including individuals from sky island populations for which we had collected trait data. We then grouped individuals by their sky island population origin, and calculated mean pairwise genetic distances across all populations. We constructed UPGMA trees from these genetic distances, which we used for evolutionary models of song characteristics and beak lengths. The topologies of these population trees were similar to the topology of our ML tree. The major difference between these trees is the branching pattern at very low levels of divergence - whereas the ML tree does not have resolution among very closely related sky island populations, the population distance trees exhibit bifurcating topologies for these populations. To account for phylogenetic uncertainty, especially for topological uncertainty at these low levels of divergence but also for branch length uncertainty, we built ten bootstrap population distance trees. We then modeled trait evolution for each of these trees, and compared the results of these models to the results on the original population tree.

*Bayesian phylogenetic reconstruction and estimation of divergence times*
To estimate the divergence time for the most recent common ancestor of the EDCS species complex, we used a coalescent-based, Bayesian approach in BEAST v1.10.2. Our alignment contained 16 species as tips, with sequence data from four nuclear introns (BRM: 530 bp, CHDZ: 551 bp, MB: 806 bp, TGFb2: 641 bp) and the mtDNA genes ATP6 (675 bp) and ND2 (1041 bp), which were treated as a single locus by concatenation. We used GTR+G substitution models for all loci, with priors on rates based on substitution rate estimates for Hawaiian honeycreepers (oscine songbirds, like sunbirds) from Lerner et al. 2011(Lerner et al. 2011). These priors were specified as normal distributions (mean ± SD: TGFb2, .0017 ± 2.2 x $10^{-4}$; all other nuclear introns, .001 ± 2.8 x $10^{-4}$; mtDNA, .0275 ± .0028). We dated the most recent common ancestor of the EDCS complex by placing a normal prior age distribution with a mean of 18 My on the split between sunbirds and flowerpeckers (three of which are included in the analysis), following (Oliveros et al. 2019). To reflect the substantial uncertainty in the age of this split, we set the standard deviation of the prior distribution at 5 My, such that 95% of the prior distribution on the node age falls between 8.2 and 27.8 My.

We ran three replicate MCMC chains in BEAST, for 25 million generations each, totaling 75 million generations. We checked each for convergence in Tracer v.1.7.1 (Rambaut et al. 2018) before combining chains using Log Combiner 1.10.2. We present the Maximum Clade Credibility tree from this analysis (Figure A3).

**Figure A1**: Bivariate plots of song measurements at the individual level, with molecular lineage coded by color and shape. Molecular lineages coded as follows: Blue circles = *fuelleborni*, red squares = *loveridgei*, yellow squares = *moreaui*, purple crosses = northern (Kenyan) populations of *mediocris*, green triangles = southern (Tanzanian) populations of *mediocris*, light blue x's = *usambarica*. Variables: ind.median.gap.after = median duration of pauses between notes (s), ind.co.var.gap.after = coefficient of variation of pause durations, ind.median.peak.frequency = median peak frequency (Hz), ind.co.var.peak.frequency = coefficient of variation in peak frequency, log.ind.elements = log of the number of elements, log.duration = log duration.

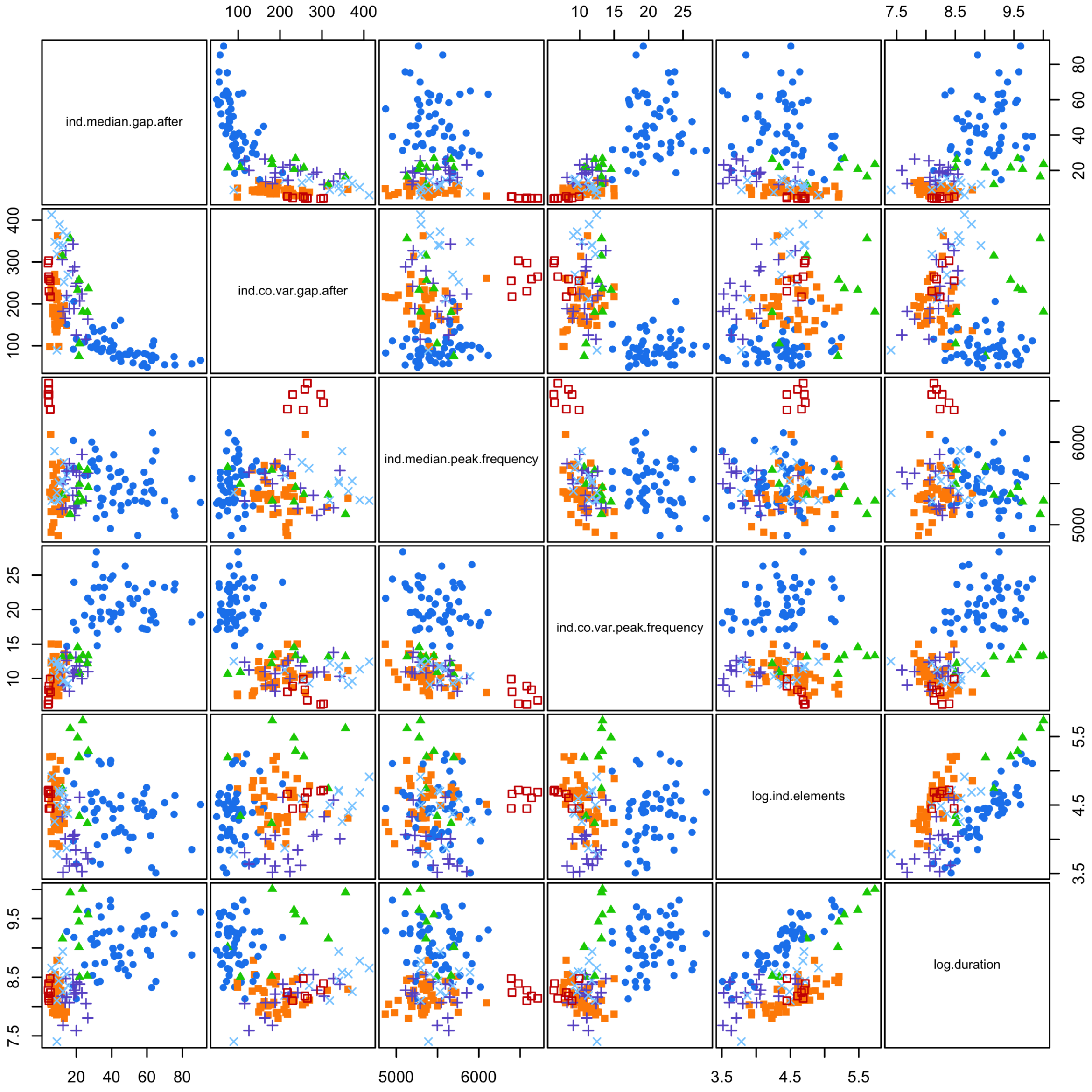

**Figure A2**. Phylogenetic analysis of the Eastern Double-collared Sunbird species complex, with outgroups. Bootstrap values are shown in blue for important nodes.

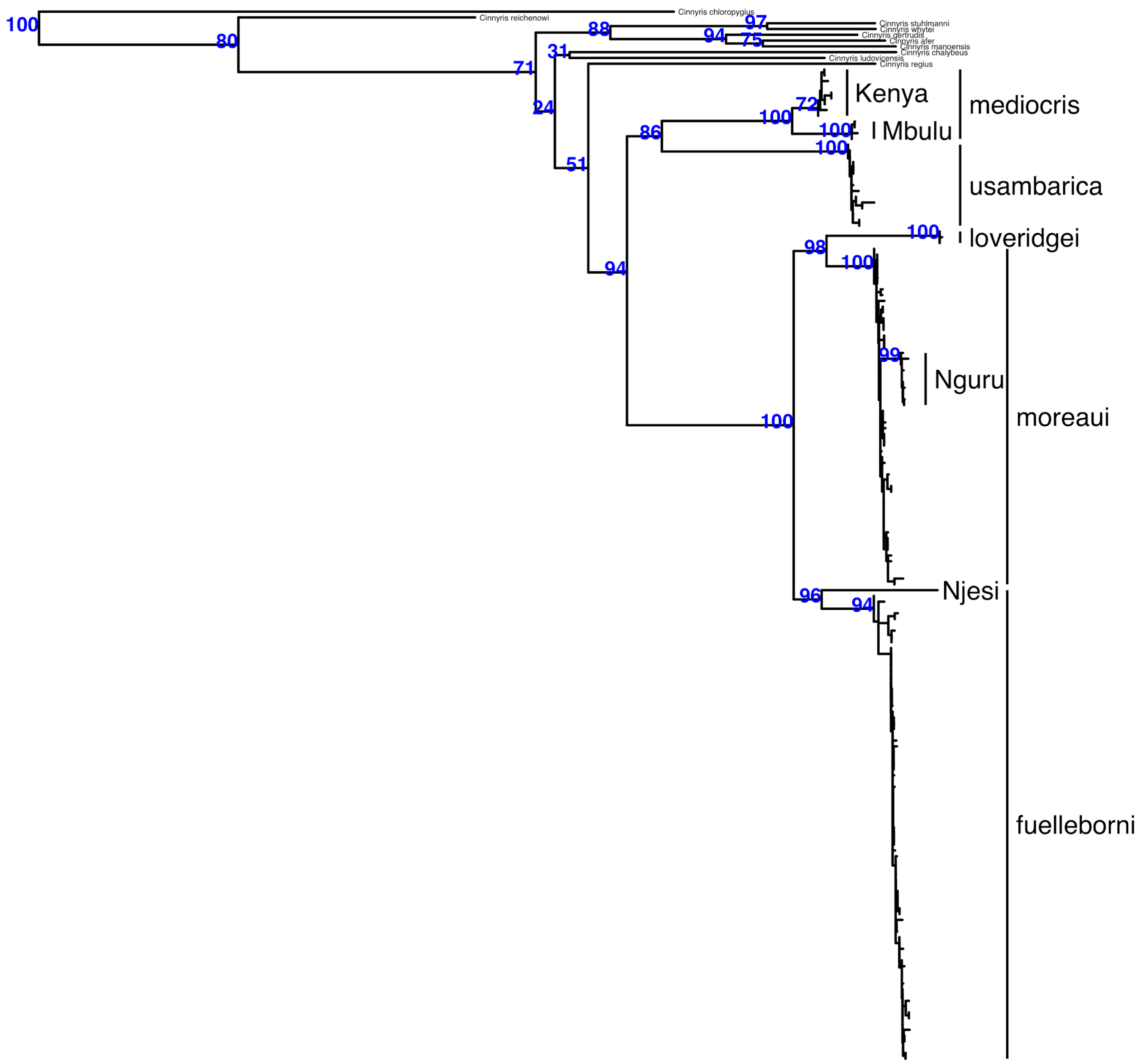

**Figure A3**. Chronogram from a Bayesian analysis of four nuclear introns and two mtDNA genes, using BEAST. Mean node age over the posterior set of trees is shown at each node, with the 95% HPD Interval indicated with a blue bar. The scale below the tree is shown in millions of years before present. The clade of interest in this study includes *Cinnyris mediocris*, *C. usambaricus*, *C. fuelleborni*, *C. loveridgei*, and *C. moreaui*.

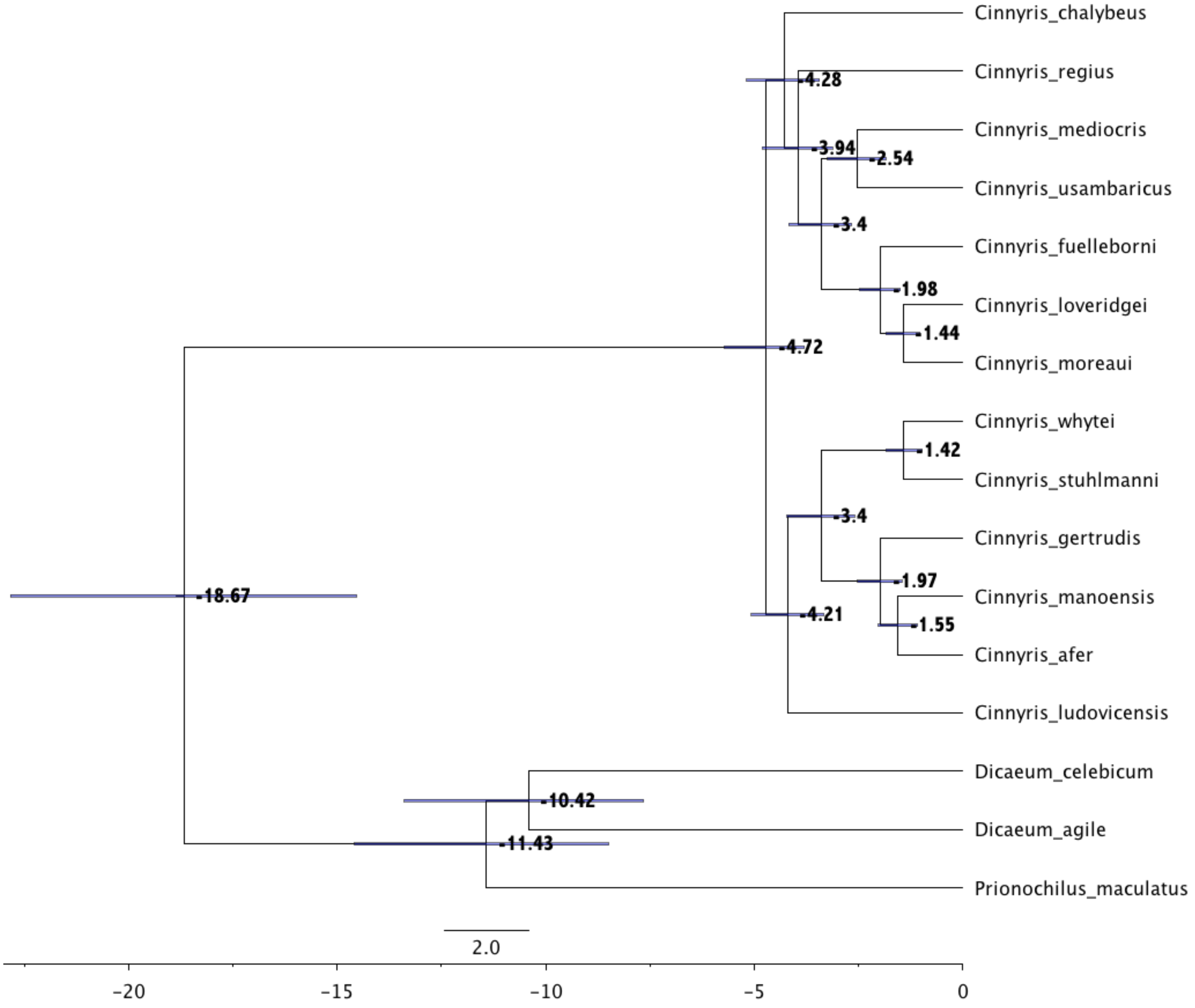

**Figures A4 - A9**. Pulse localization for the evolution of different song traits. Blue diamonds are found on branches where pulse localizations had support values > 0.2, with diamond size reflecting support, and support value shown above. Asterisks signify pulses that occur in the pulse configuration with the minimum AICc value. Sizes of gray circles correspond to mean phenotype values at tips, which each represent a geographically discrete sky island population. Species epithets are indicated at far right.

**Figure A4**.

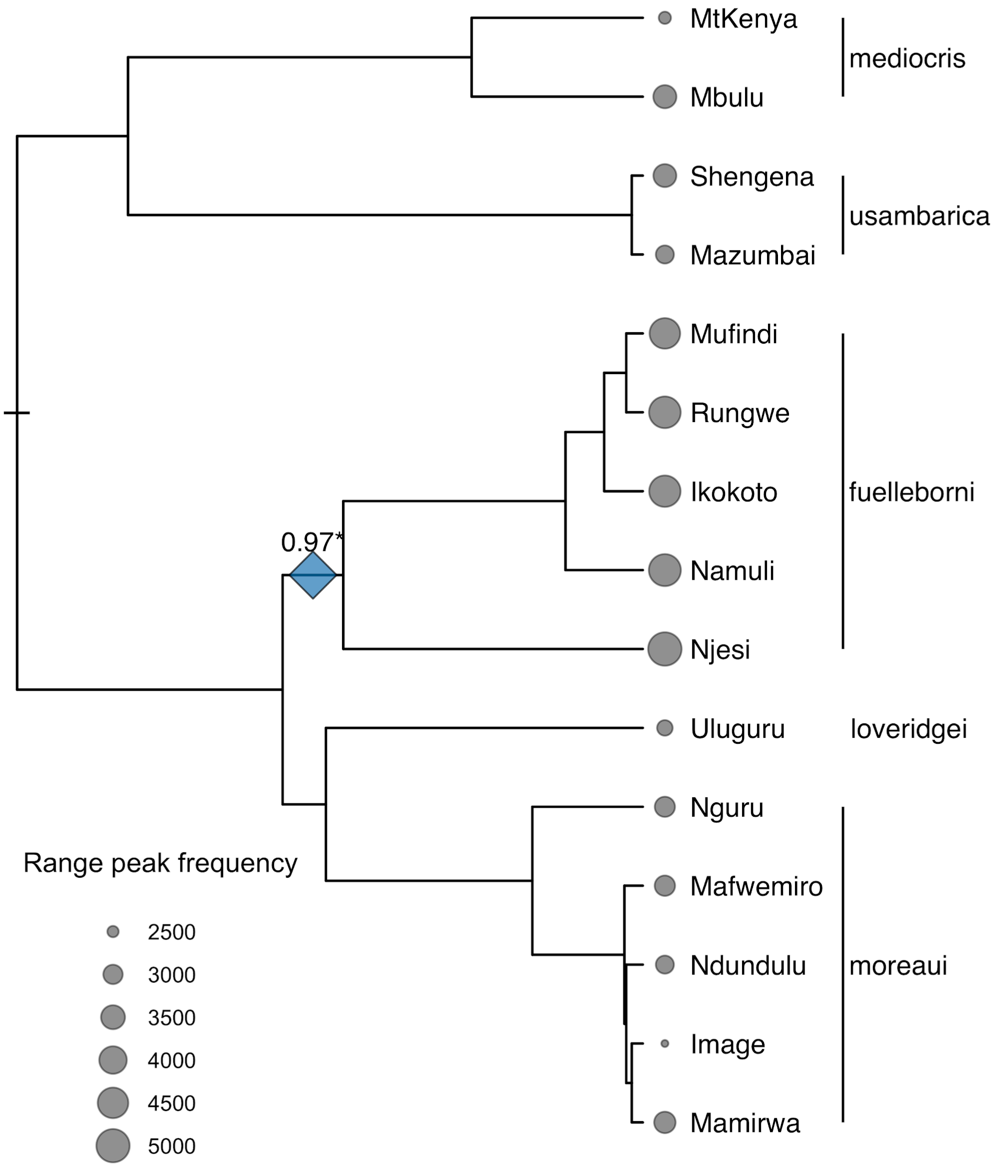

**Figure A5.**

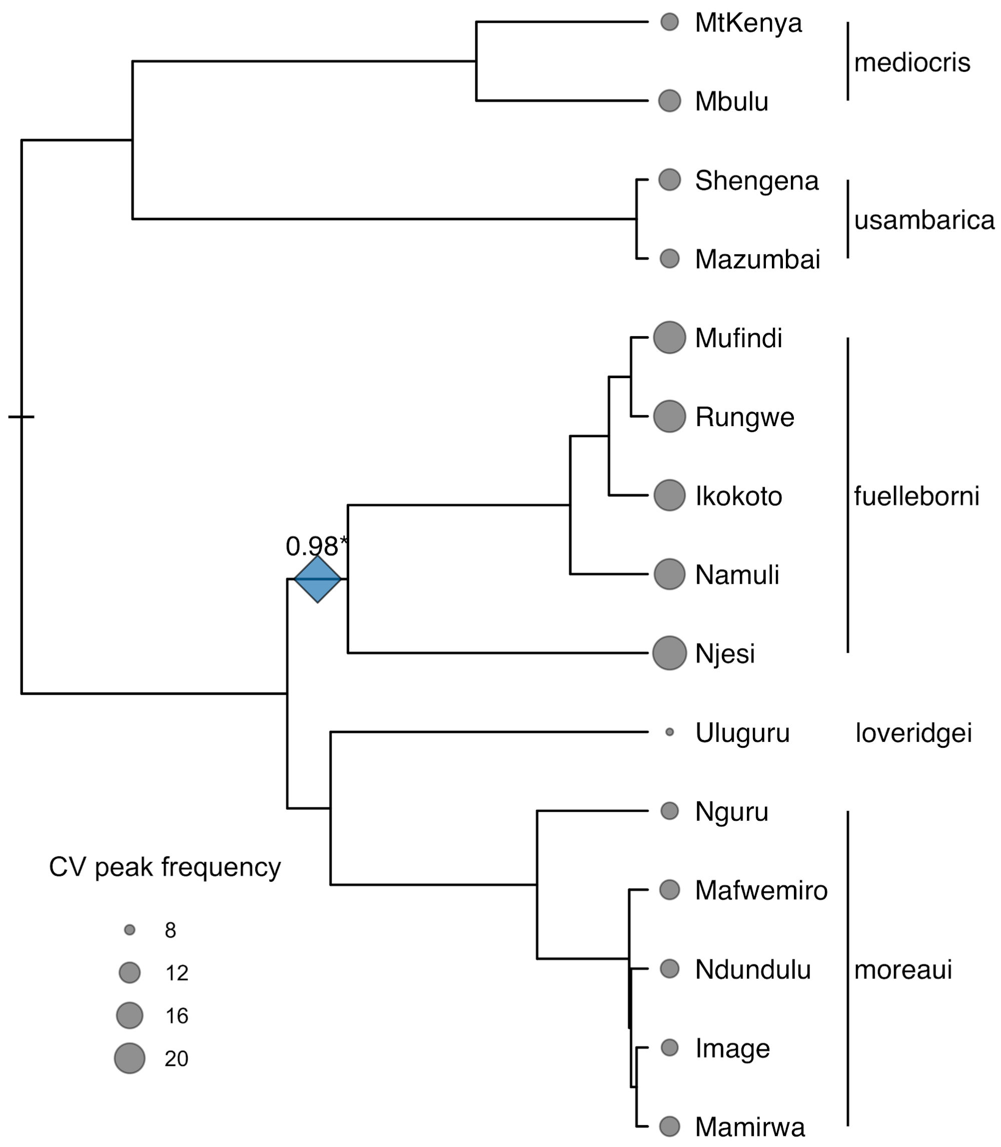
MtKenya
Mbulu
mediocris
Shengena
Mazumbai
usambarica
Mufindi
Rungwe
Ikokoto
Namuli
Njesi
fuelleborni
0.98*
Uluguru
loveridgei
Nguru
Mafwemiro
Ndundulu
Image
Mamirwa
moreaui
CV peak frequency
8
12
16
20

**Figure A6.**

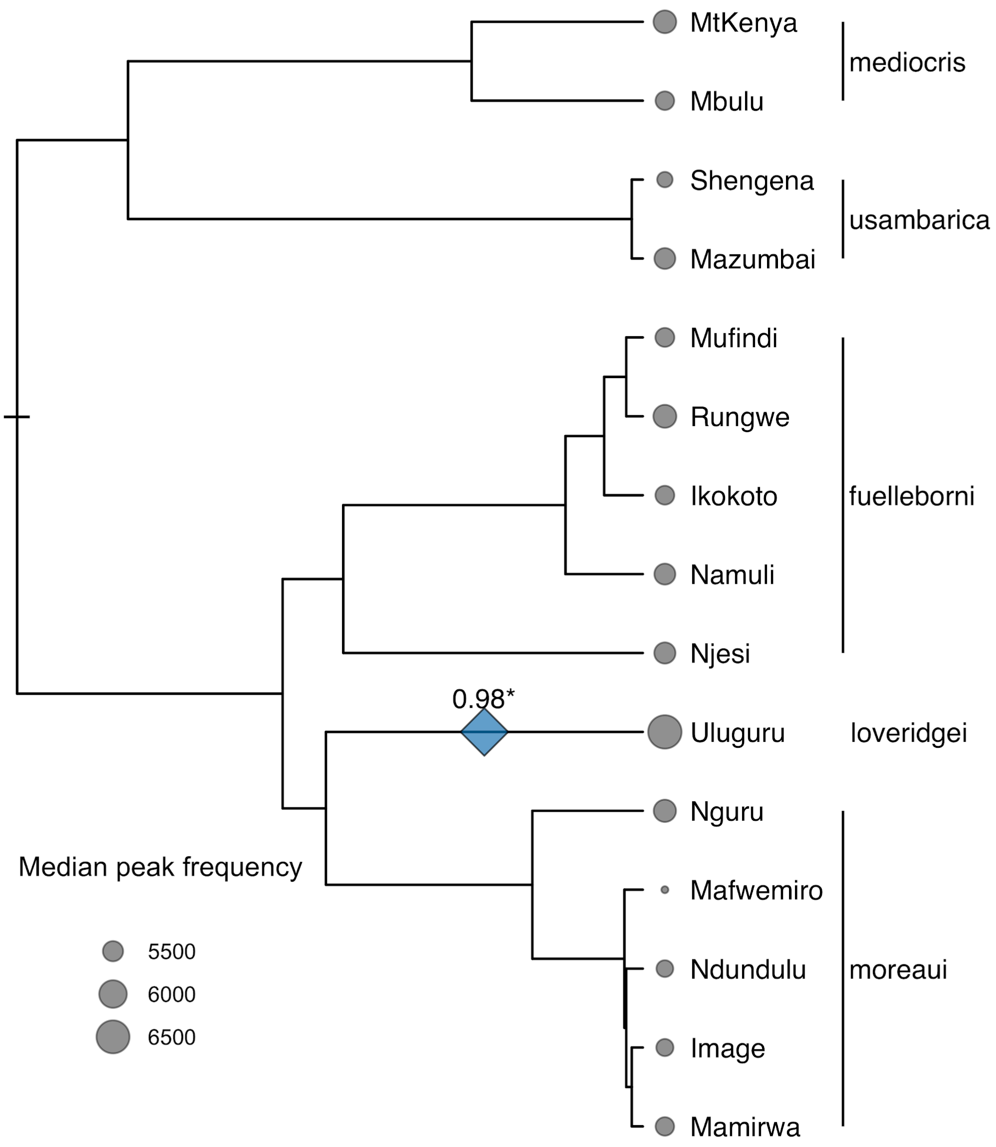

MtKenya
Mbulu
mediocris
Shengena
Mazumbai
usambarica
Mufindi
Rungwe
Ikokoto
Namuli
Njesi
fuelleborni
0.98*
Uluguru
loveridgei
Nguru
Mafwemiro
Ndundulu
Image
Mamirwa
moreaui
Median peak frequency
5500
6000
6500

**Figure A7.**

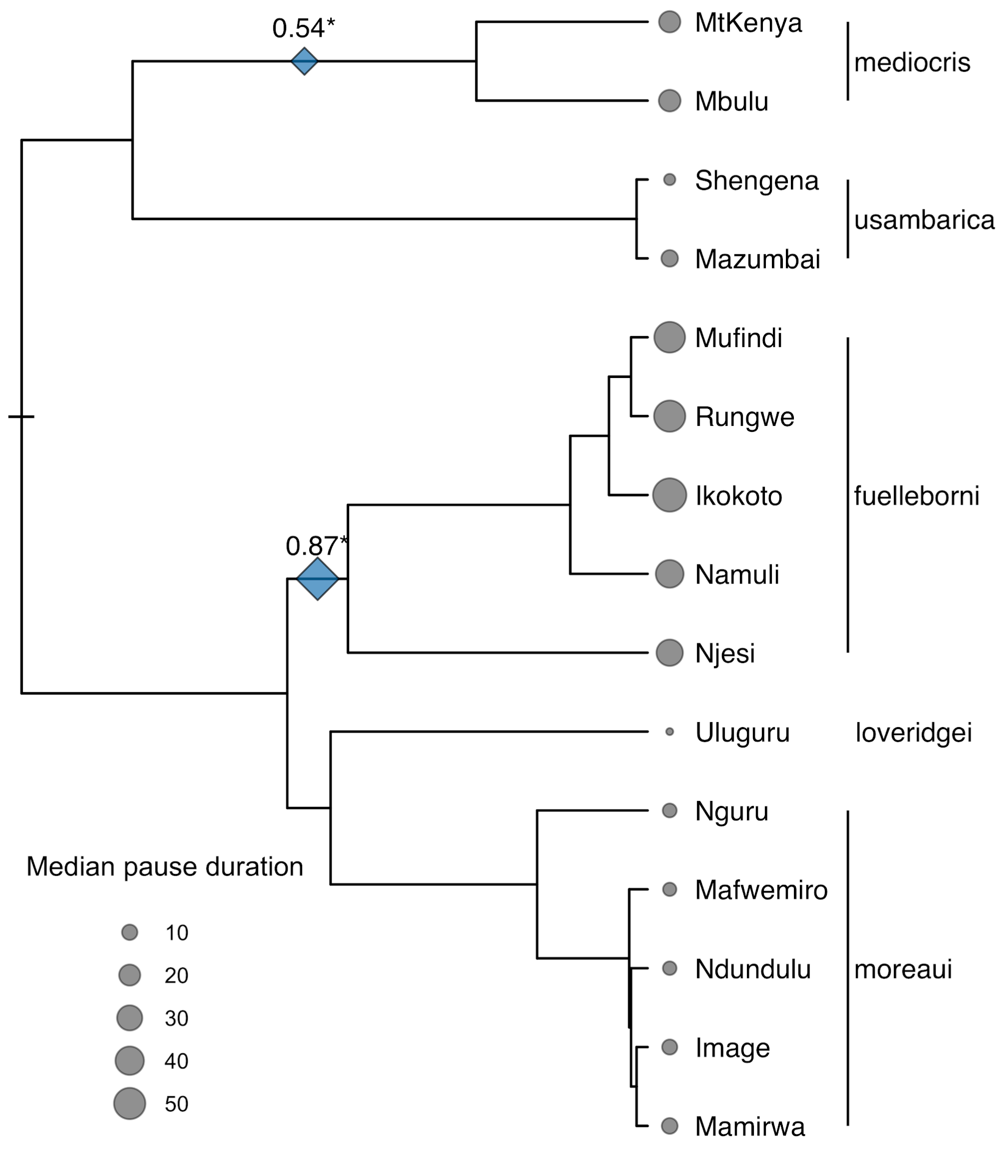
0.54*
MtKenya
Mbulu
mediocris
Shengena
Mazumbai
usambarica
Mufindi
Rungwe
Ikokoto
fuelleborni
0.87*
Namuli
Njesi
Uluguru
loveridgei
Nguru
Median pause duration
10
20
30
40
50
Mafwemiro
Ndundulu
moreaui
Image
Mamirwa

**Figure A8.**

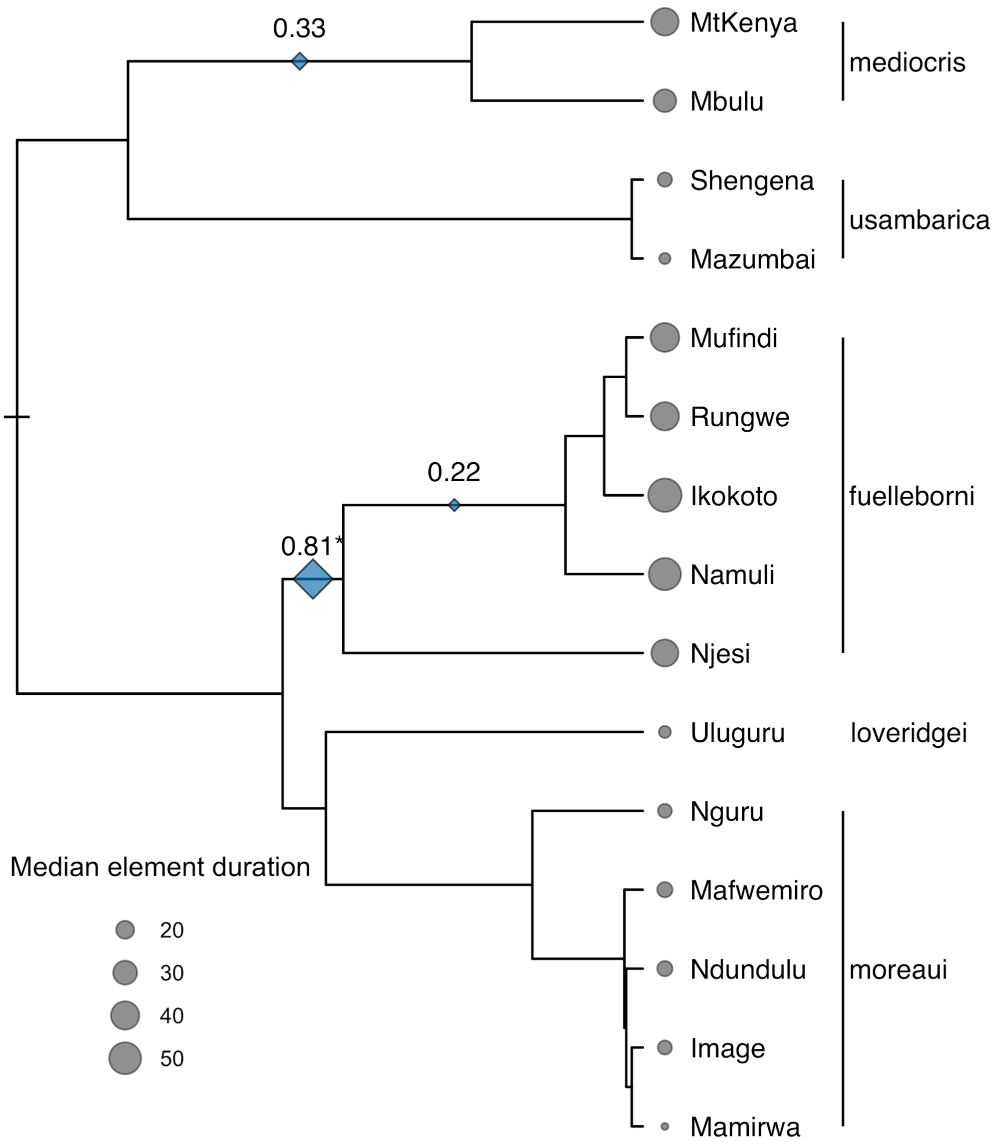

0.33
0.22
0.81*
MtKenya
Mbulu
Shengena
Mazumbai
Mufindi
Rungwe
Ikokoto
Namuli
Njesi
Uluguru
Nguru
Mafwemiro
Ndundulu
Image
Mamirwa
mediocris
usambarica
fuelleborni
loveridgei
moreaui
Median element duration
20
30
40
50

**Figure A9.**

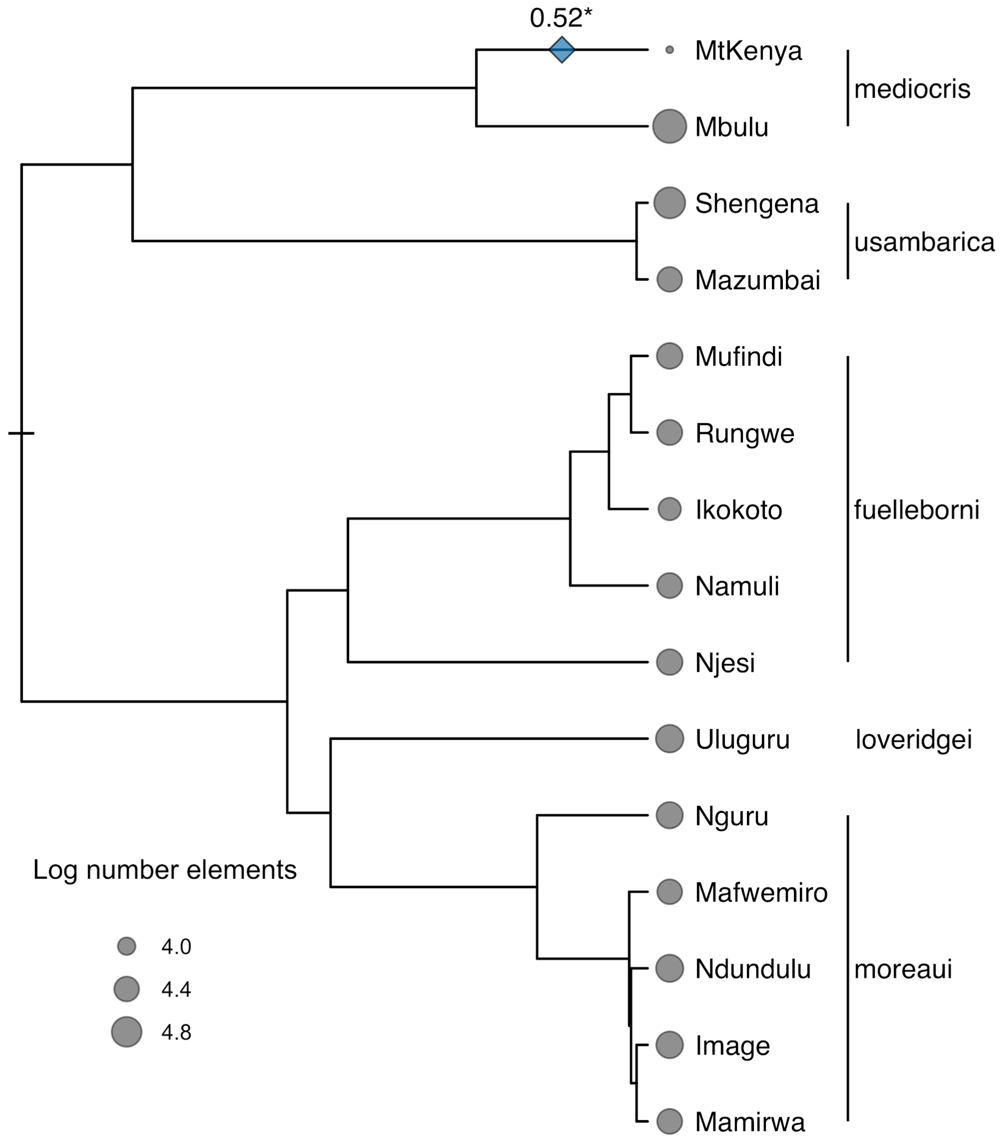
0.52*
MtKenya
Mbulu
mediocris
Shengena
Mazumbai
usambarica
Mufindi
Rungwe
Ikokoto
Namuli
Njesi
fuelleborni
Uluguru
loveridgei
Nguru
Mafwemiro
Ndundulu
Image
Mamirwa
moreaui
Log number elements
4.0
4.4
4.8

### References


Backström, N., N. Karaiskou, E. H. Leder, L. Gustafsson, C. R. Primmer, A. Qvarnström, and H. Ellegren. 2008. A gene-based genetic linkage map of the collared flycatcher (Ficedula albicollis) reveals extensive synteny and gene-order conservation during 100 million years of avian evolution. Genetics 179:1479–1495.

Burnham, K. P., and D. R. Anderson. 2003. Model Selection and Multimodel Inference: A Practical Information-Theoretic Approach. Springer Science & Business Media.

Chesser, R. T. 1999. Molecular Systematics of the Rhinocryptid Genus Pteroptochos. The Condor 101:439–446.

Griffiths, R., and R. M. Korn. 1997. A CHD1 gene is Z chromosome linked in the chicken Gallus domesticus. Gene 197:225–229.

Hunt, J. S., E. Bermingham, and R. E. Ricklefs. 2001. Molecular Systematics and Biogeography of Antillean Thrashers, Tremblers, and Mockingbirds (Aves: Mimidae). The Auk 118:35–55.

Katoh, K., K. Kuma, H. Toh, and T. Miyata. 2005. MAFFT version 5: improvement in accuracy of multiple sequence alignment. Nucleic acids research 33:511–518.

Kimball, R. T., E. L. Braun, F. K. Barker, R. C. K. Bowie, M. J. Braun, J. L. Chojnowski, S. J. Hackett, et al. 2009. A well-tested set of primers to amplify regions spread across the avian genome. Molecular phylogenetics and evolution 50:654–660.

Lanfear, R., B. Calcott, S. Y. W. Ho, and S. Guindon. 2012. PartitionFinder: combined selection of partitioning schemes and substitution models for phylogenetic analyses. Molecular biology and evolution 29:1695–1701.

Lanfear, R., P. B. Frandsen, A. M. Wright, T. Senfeld, and B. Calcott. 2017. PartitionFinder 2: New Methods for Selecting Partitioned Models of Evolution for Molecular and Morphological Phylogenetic Analyses. Molecular biology and evolution 34:772–773.

Lerner, H. R. L., M. Meyer, H. F. James, M. Hofreiter, and R. C. Fleischer. 2011. Multilocus resolution of phylogeny and timescale in the extant adaptive radiation of Hawaiian honeycreepers. Current biology: CB 21:1838–1844.

Miller, M. A., W. Pfeiffer, and T. Schwartz. 2010. Creating the CIPRES Science Gateway for inference of large phylogenetic trees. Pages 1–8 *in*2010 Gateway Computing Environments Workshop (GCE). ieeexplore.ieee.org.

Oliveros, C. H., D. J. Field, D. T. Ksepka, F. K. Barker, A. Aleixo, M. J. Andersen, P. Alström, et al. 2019. Earth history and the passerine superradiation. Proceedings of the National Academy of Sciences of the United States of America.

Rambaut, A., A. J. Drummond, D. Xie, G. Baele, and M. A. Suchard. 2018. Posterior Summarization in Bayesian Phylogenetics Using Tracer 1.7. Systematic biology 67:901–904.

Sorenson, M. D., J. C. Ast, D. E. Dimcheff, T. Yuri, and D. P. Mindell. 1999. Primers for a PCR-based approach to mitochondrial genome sequencing in birds and other vertebrates. Molecular phylogenetics and evolution 12:105–114.

Stamatakis, A. 2014. RAxML version 8: a tool for phylogenetic analysis and post-analysis of large phylogenies. Bioinformatics 30:1312–1313.
