## Supplementary Information 1 for "Inferring punctuated evolution in the learned songs of African sunbirds"

### Supplementary Information for: Inferring punctuated evolution in the learned songs of African sunbirds

#### 1 Introduction

##### 1.1 Overview

We consider two models for how a trait's mean evolves over time: pulsed evolution, and Brownian motion (BM). To fit the first model, we iterate over all possible placements of up to 6 pulses on the tree. For each pulse configuration, we find the maximum likelihood estimates for the ancestral mean, variance of pulse displacements, and variance of short-term bounded evolution. We then use the corrected Akaike information criterion (AICc) of each pulse configuration to choose the likeliest pulse configuration, and use the AICc weights of each configuration to estimate the values of the parameters under the pulse model. To fit the BM model, we find the maximum likelihood estimates (MLEs) of the ancestral mean, strength of BM, and variance of short-term bounded evolution.

Model selection is accomplished via the comparison of AICc scores. Because phylogenetically correlated data does not satisfy the assumption of independence that is used in the derivation of the AICc score as the Kullback–Leibler divergence (KL) between the empirical distribution of the data and the model under consideration [1, 10, 7, 8], standard approaches to model selection using AICc differences may not be appropriate [3]. To understand our ability to recover evidence for either model using AICc in the presence of phylogenetic correlations and measurement errors, we performed a set of simulation studies.

We remark that under the above pulse models, we do not model the dynamics (i.e. rates) of the pulses. Instead, we only attempt to place them on the evolutionary tree.

##### 1.2 Probability density of mean traits

For the phylogeny considered in this work, denote by $\mu$ the average trait measure of the common ancestor of all the present-day populations, which have mean trait values $Z_1, \dots, Z_N$, where $N$ is the number of populations. Denote a shift along the $k$th edge in the tree by $X_k$, where $1 \leq k \leq 2(N-1)$.

Assume that the trait shifts are independent and each is distributed as a normal distribution with mean 0 and variance $d_k^2$. Similarly, assume the trait of the $i$th (present-day) measured trait is normally-distributed and has variance $\sigma_p^2 + D_i^2$, where $\sigma_p^2$ denotes the strength of the white noise fluctuations in the mean trait (i.e. non-accumulating contributions to the mean trait throughout the species' lineage, which amount to bounded evolution), and $D_i^2$ is the variance of the measurement error. It follows that the probability density function of the vector $Z = (Z_1, \ldots, Z_N)^T$ is

$$g_Z(z_1, \ldots, z_N) = \frac{1}{\sqrt{(2\pi)^N |\Sigma|}} e^{-(Z-\mu_0)^T \Sigma^{-1} (Z-\mu_0)}, \tag{1}$$

where $\mu_0 = (\mu, \ldots, \mu)^T$ is the mean vector, and $\Sigma$ is the covariance matrix given by

$$\Sigma_{ii} = \sigma_p^2 + D_i^2 + \sum_{k \in \ell_i} d_k^2, \tag{2}$$

$$\Sigma_{ij} = \sum_{k \in \ell_i \cap \ell_j} d_k^2 \quad (i \neq j), \tag{3}$$

where $\ell_i$ is the set of indices of the edges in the lineage of the $i$th population, and $\ell_i \cap \ell_j$ is the set of indices of the edges in the common lineage of the $i$th and $j$th populations.

##### 1.3 Choice of displacement variance

We consider two models for the trait displacement along an edge:

1. A Brownian motion model, in which the trait evolves continuously along the edge as BM, so that

$$X_k \sim N(0, \sigma_{bm}^2 \tau_k), \tag{4}$$

   where $\tau_k$ is the branch length, and $d_k = \sigma_{bm}^2 \tau_k$.

2. A pulse model, in which the $k$th edge has $n_k$ prescribed displacement pulses (possibly zero), and is therefore distributed according to the sum of $n_k$ independent and identical normally-distributed random variables, so that

$$X_k \sim N(0, n_k \sigma_d^2) \tag{5}$$

   and $d_k = n_k \sigma_d^2$.

#### 2 Model fitting

##### 2.1 Data rescaling and phylogeny storage

Each data set is shifted by its mean, and then rescaled by the difference between the maximum and minimum of the data set. This is done to make the below-described methods computationally stable.

Handling and visualization of the phylogenetic tree is accomplished using the ETE Toolkit [6] in Python.

##### 2.2 Computing the likelihood for a fixed covariance matrix

For a vector of measurements $Z$ and a covariance matrix $\Sigma$ as given above, we can quickly compute the likelihood using Felsenstein's independent contrasts method [4, 5], which uses independent contrasts to compute both the likelihood evaluated at the MLE for $\mu$, and the MLE for $\mu$ conditioned on the covariance matrix. Thus, we assume that $\mu$ is a known function of the variance parameters.

We remark that although the method was introduced specifically for the BM model, it is equally applicable to any multivariate normal model with a covariance matrix which corresponds to summing variances across the shared lineages of any two tips.

##### 2.3 Fitting MLE parameters

Generally, the likelihood functions for different (even normalized) data sets are very different, and very flat—except when very close to the maximum. Both models require good initial guesses for a gradient descent algorithm to successfully converge. Below we describe how to get such guesses.

##### 2.3.1 Fitting the BM model

For the BM model, denote the negative logarithm of the likelihood

$$l(\sigma_p^2, \sigma_{bm}^2) = -\ln L(\sigma_p^2, \sigma_{bm}^2, \mu(\sigma_p^2, \sigma_{bm}^2)), \tag{6}$$

where $L$ is the likelihood, and $\mu(\sigma_p^2, \sigma_{bm}^2)$ is co-computed using the independent contrasts method. We evaluate $l(\sigma_p^2, \sigma_{bm}^2)$ on a $180 \times 180$ uniform mesh over $[10^{-30}, 1] \times [0, 4]$, and use the minimum as an initial guess for finding the minimum of $l$ via gradient descent.

##### 2.3.2 Fitting the pulse model

**Parameter values** Consider a fixed configuration of pulses. The pulses divide the tip samples into several regimes, so that all tip samples within a regime share the same history of pulses. Within the $k$th regime, the likelihood of $\sigma_p^2$ is given by

$$L_k(\sigma_p^2) = \prod_{i \in K} \frac{1}{\sqrt{2\pi(\sigma_p^2 + D_i^2)}} e^{-(x_i - \mu_k(\sigma_p^2))^2/2(\sigma_p^2 + D_i^2)}, \tag{7}$$

where $\mu_k(\sigma_p^2)$ is obtained by inverse-variance weighing. To estimate $\sigma_p^2$, we maximize $G(\sigma_p^2) = \prod_k g_k(\sigma_p^2)$ to obtain the estimate $\tilde{\sigma_p}^2$ .

Having obtained $\tilde{\sigma}_p^2$, we minimize the (now univariate) conditioned negative log-likelihood function $h(\sigma_d^2) = l(\hat{\mu}(\tilde{\sigma}_p^2, \sigma_d^2), \tilde{\sigma}_p^2, \sigma_d^2)$ to obtain the estimate $\tilde{\sigma}_d^2$.

Finally, we minimize $l$ using gradient descent, using $(\tilde{\sigma}_p^2, \tilde{\sigma}_d^2)$ as an initial guess. In practice, this initial guess is typically very close to the true minimum, and the gradient descent algorithm converges after a very small number of iterations.

**Importance of incorporating measurement errors in initial guess** We note that when the measurement errors are zero, i.e. $D_i = 0$ for all $i$, (7) may be solved analytically. This is therefore a more efficient way for obtaining an estimate for $\sigma_p^2$ when the the measurement errors are small, or are ignored. However, such an approximation is not sufficient when the measurement errors are of the magnitude exhibited in this study; in this case, the likelihood function (which takes into account measurement errors) is very flat near the estimated $\sigma_p^2$, as illustrated in Figure S1.

**Finding likely pulse configurations** We repeat the above procedure for each possible configuration of up to 4 pulses, and choose the configuration with the smallest AICc score. This can be visualized as solving the minimization problem for different matrix shapes, the first few of which are illustrated in Figure S2.

For each dataset we considered, no more than 2 pulses were chosen. For the purposes of AICc score computation, we count each pulse as an additional parameter.

We found that using an MCMC-type algorithm with cached AICc scores for each pulse configuration allows for exploring many more pulse configurations. However, as we do not expect many pulses in the present datasets, we used the faster (due to the relatively small sample space) brute-force calculation approach.

#### 2.4 Results of model fitting

Using the algorithm described above, we fit the model parameters for each trait. The results for the BM and pulse models are given in Table S2 and Table S2, respectively.

### 3 Interpretation of ΔAICc for each trait

#### 3.1 Phylogenetic non-independence and use of AICc values

The AICc value of a model with maximum likelihood $L$, $k$ parameters, and a sample size $n$ is given by

$$\text{AICc} = 2k - 2\log L + \frac{2k(k+1)}{n-k-1}. \tag{8}$$

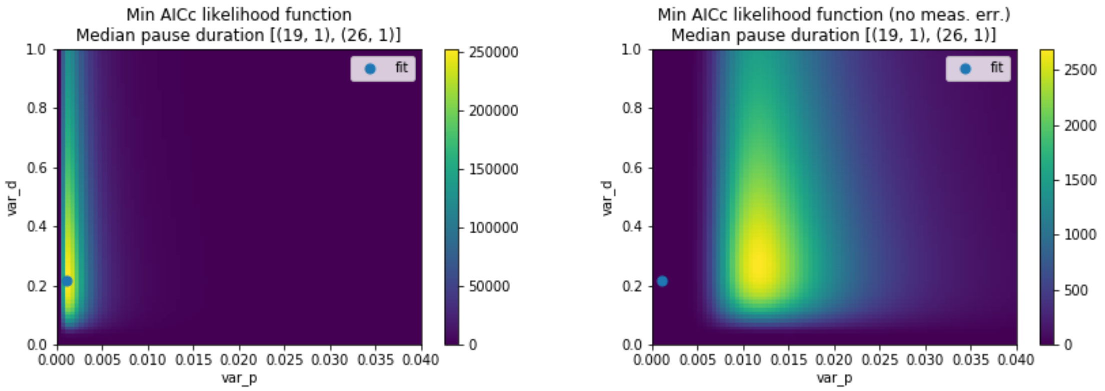


Figure S1: The likelihood function for an optimal configuration of pulses, with and without measurement errors. Although estimating $\sigma_p^2$ analytically is possible in the in the latter case, this estimate is far away from the $\sigma_p^2$ value in the former case. When this analytic approximation is used as an initial guess for fitting the parameters in a model which incorporates measurement errors, a standard gradient descent algorithm typically does not converge to the true maximum, due to the flatness of the likelihood surface. The small disc on each graph denotes the true maximum for the model with measurement errors. The shift of the maximum to the right when measurement errors are ignored is evident in all the traits considered, and suggests that the true value of $\sigma_p^2$ is usually finite and small, but hidden by the measurement errors.

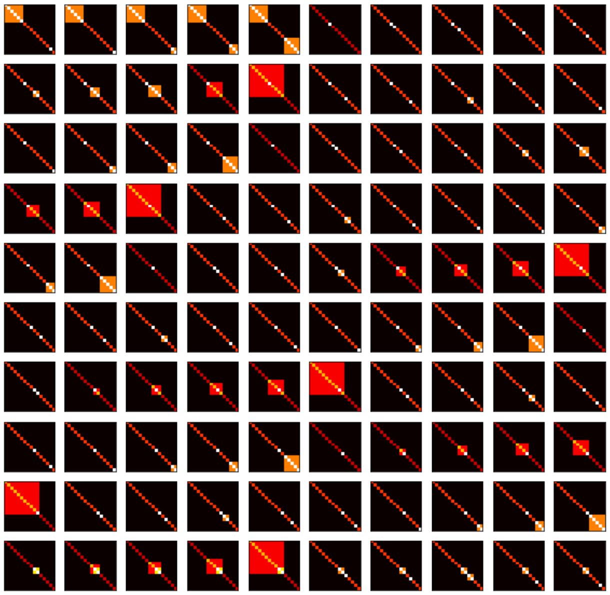

Figure S2: The first 100 matrix shapes iterated over for the pulse model. Each color corresponds to a different constant. Iterating over up to four pulses corresponds to iterating over 35,960 possible configurations.

| Trait | BM $\sigma_p^2$ | BM $\sigma_{bm}^2$ | BM $\mu$ | $\text{AICc}_{\text{BM}}$ |
|---|---|---|---|---|
| CV frequency bandwidth | 6.31E-27 | 1.02E+04 | 8.29E+01 | 142.918 |
| CV frequency change | 9.36E-28 | 1.49E+03 | 7.78E+01 | 117.882 |
| max peak frequency (Hz) | 2.95E-24 | 5.71E+06 | 7.15E+03 | 230.054 |
| log duration | 1.77E-30 | 3.53E+00 | 8.56E+00 | 18.151 |
| min peak frequency (Hz) | 4.40E+04 | 5.59E+06 | 3.63E+03 | 240.074 |
| log median frequency change | 2.77E-31 | 0.00E+00 | -2.75E+00 | -4.333 |
| median pause duration | 2.86E-27 | 2.84E+03 | 1.81E+01 | 116.805 |
| median element duration | 1.80E-27 | 3.01E+03 | 2.48E+01 | 118.283 |
| range peak frequency | 7.26E-24 | 1.41E+07 | 3.41E+03 | 244.322 |
| CV pause duration | 1.12E-25 | 8.77E+04 | 2.13E+02 | 171.117 |
| CV peak frequency | 2.69E-28 | 4.56E+02 | 1.30E+01 | 85.409 |
| number of elements | 1.85E-30 | 0.00E+00 | 4.45E+00 | 11.147 |
| log median bandwidth | 3.67E-30 | 3.66E+00 | 6.22E+00 | 28.168 |
| median peak frequency | 1.98E-24 | 3.68E+06 | 5.62E+03 | 222.583 |

Table S2: Fit values for the BM model.

| Trait | Pulse $\sigma_p^2$ | Pulse $\sigma_d^2$ | Pulse $\mu$ | Pulse positions | $\text{AICc}_{\text{PE}}$ | $\Delta A$ |
|---|---|---|---|---|---|---|
| CV frequency bandwidth | 2.00E+01 | 1.86E+03 | 1.02E+02 | [(9, 1), (26, 1)] | 144.090 | -1.173 |
| CV frequency change | 5.81E+00 | 2.03E+02 | 8.26E+01 | [(19, 1)] | 117.116 | 0.766 |
| max peak frequency (Hz) | 1.72E+01 | 1.06E+06 | 7.68E+03 | [(8, 1), (27, 1)] | 226.736 | 3.318 |
| log duration | 1.11E-05 | 9.36E-01 | 8.15E+00 | [(19, 1)] | 12.514 | 5.637 |
| min peak frequency (Hz) | 5.79E-14 | 6.93E+05 | 3.86E+03 | [(5, 1), (19, 1)] | 237.322 | 2.753 |
| log median frequency change | 1.65E-06 | 1.11E+00 | -2.75E+00 | [] | -4.331 | -0.001 |
| median pause duration | 3.20E+00 | 6.14E+02 | 7.91E+00 | [(19, 1), (26, 1)] | 111.195 | 5.610 |
| median element duration | 1.04E-02 | 7.86E+02 | 1.65E+01 | [(19, 1)] | 113.136 | 5.147 |
| range peak frequency | 2.09E+02 | 3.13E+06 | 2.97E+03 | [(19, 1)] | 232.711 | 11.610 |
| CV pause duration | 6.66E-01 | 1.67E+04 | 2.18E+02 | [(19, 1)] | 170.324 | 0.793 |
| CV peak frequency | 1.93E-01 | 1.04E+02 | 1.06E+01 | [(19, 1)] | 71.645 | 13.764 |
| number of elements | 1.10E-05 | 4.52E-01 | 4.48E+00 | [(25, 1)] | 7.516 | 3.630 |
| log median bandwidth | 2.37E-05 | 9.20E-01 | 6.18E+00 | [(21, 1), (26, 1)] | 26.383 | 1.785 |
| median peak frequency | 1.15E+01 | 1.19E+06 | 5.44E+03 | [(9, 1)] | 210.954 | 11.629 |

Table S2: Fit values for the pulse model, as well as the $\Delta A$ values. Pulse positions are given a sequence of edge numbers, and the number of pulses on them (e.g. $(19, 1)$ refers to one pulse on the 19th edge). The edge numbers are given in Figure S8.

This value is an approximation of the cross entropy of the data's empirical distribution, and a model under consideration. For phylogenetically correlated data, the empirical distribution of the data may not be a good approximation of the true distribution. It therefore follows that in the context of model selection, ΔAICc values may not be interpretable in the usual way [3]. For model selection purposes in the context of this work, we empirically verify that reliable model selection with correlated data may be accomplished by specifying a different cutoff for interpreting a ΔAICc value as strong evidence, which depends on the model parameters.

Lastly, we note that the likelihood function for the pulse model is the likelihood conditioned on some specific outcome of pulse dynamics (e.g. the pulse normal model in [9]). The stricter ΔAICc score threshold also compensates for the pulse model's inherently greater likelihood.

#### 3.2 Simulating data to generate distributions of ΔAICc values

##### 3.2.1 Overview

For each song variable, we generated 500 simulated datasets using parameters close to the MLE parameters for each of the two models of interest. We also simulated measurement error for these data sets based on the measurement errors from our empirical data sets. We then fit our two models of interest to these simulated data sets, and present the distribution of differences between the BM and pulsed evolution AICc scores that we recovered,

$$\Delta A = \mathrm{AICc}_{\mathrm{BM}} - \mathrm{AICc}_{\mathrm{PE}}. \tag{9}$$

Examining these distributions allows us to identify where relative support for BM or pulsed evolution is strongest.

##### 3.2.2 Simulated datasets

For each real dataset with MLE parameters, we generate 500 BM-generated datasets, and 500 pulse-generated datasets. Each dataset has artificial measurement errors similar in magnitude to those in the real datasets.

**BM-generated datasets** We perturb each parameter by up to 5% uniformly, and sample once from the resulting distribution to obtain the true trait mean $x_i$ at each leaf. At the $i$th leaf, we choose an artificial measurement error variance $e_i^2 = \alpha_i^2 x_i^2$ with $\alpha_i$ uniformly chosen from $(0, 0.3)$, and sample from $N(x_i, e_i^2)$ $K_i$ times, where $K_i$ is a uniformly chosen integer between 4 and 30. The $i$th component of the simulated dataset is then the mean of these artificial samples, with the typical associated measurement errors.

**Pulse-generated datasets** We placed one or two pulses on the tree, with uniform probability of placement across the tree, and repeat the same procedure as above.

##### 3.2.3 Analyzing simulated datasets

For each trait of interest, we analyzed all 1000 corresponding simulated data sets, and separately bin the $\Delta A$ values for BM- and pulse-generated data. When fitting the pulse model, we attempt to place up to 2 pulses. The results of this procedure may be seen in Figure S3. Crucially, we see that we consistently recovered a substantial fraction of positive $\Delta A$ values (suggesting greater support for pulsed evolution) when data are generated under BM, while the reverse is not true. Raw $\Delta A$ values are biased towards greater support for pulsed evolution than BM. Thus we must be cautious in our interpretation.

##### 3.2.4 Interpretation of ΔAICc values

Given the dependence of $\Delta A$ values on model parameter values, we assessed the evidence for models individually for each trait. Informally, we consider there to be strong relative support for the pulsed evolution model when the $\Delta A$ score is greater than the 95th percentile, and greater than the 5th percentile for the pulse-generated data. For the datasets used in this work, this means the calibrated cutoff should simply be the BM-generated 95th percentile. The percentile values, as well as each trait's $\Delta A$ value, are tabulated in Table S2.

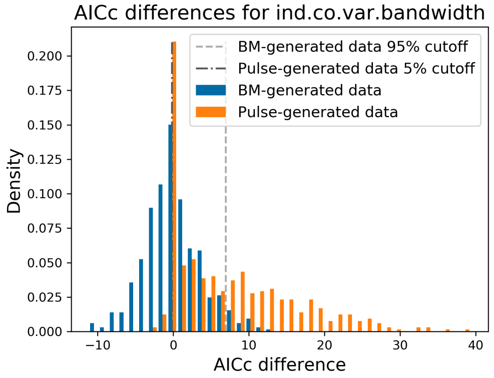
AICc differences for ind.co.var.bandwidth
BM-generated data 95% cutoff
Pulse-generated data 5% cutoff
BM-generated data
Pulse-generated data
Density
AICc difference

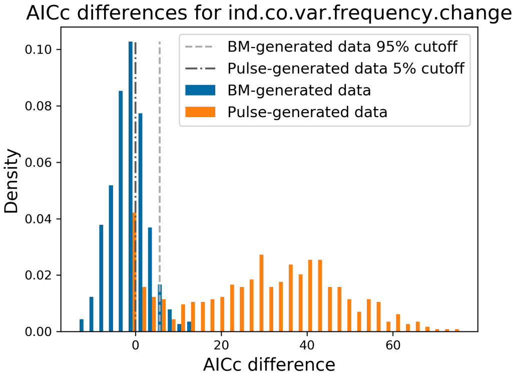
AICc differences for ind.co.var.frequency.change
BM-generated data 95% cutoff
Pulse-generated data 5% cutoff
BM-generated data
Pulse-generated data
Density
AICc difference

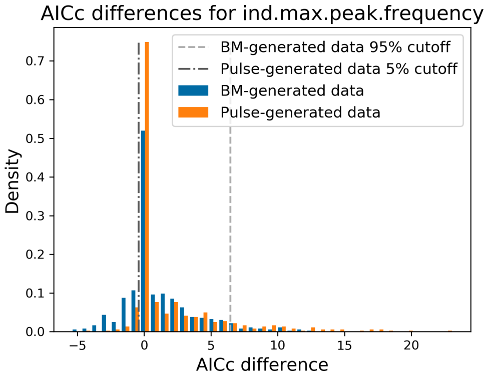
AICc differences for ind.max.peak.frequency
BM-generated data 95% cutoff
Pulse-generated data 5% cutoff
BM-generated data
Pulse-generated data
Density
AICc difference

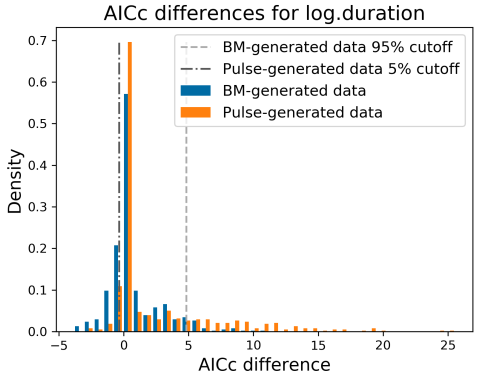
AICc differences for log.duration
BM-generated data 95% cutoff
Pulse-generated data 5% cutoff
BM-generated data
Pulse-generated data
Density
AICc difference

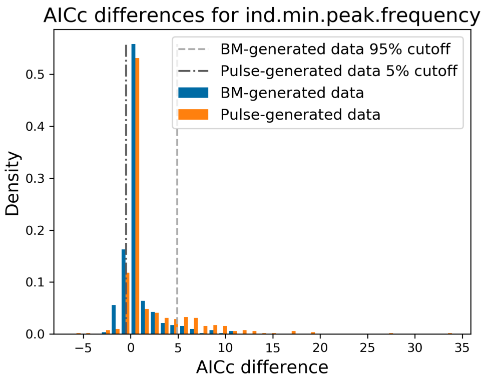
AICc differences for ind.min.peak.frequency
BM-generated data 95% cutoff
Pulse-generated data 5% cutoff
BM-generated data
Pulse-generated data
Density
AICc difference

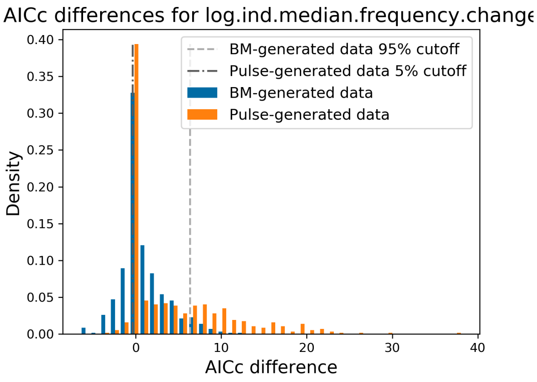
AICc differences for log.ind.median.frequency.change
BM-generated data 95% cutoff
Pulse-generated data 5% cutoff
BM-generated data
Pulse-generated data
Density
AICc difference

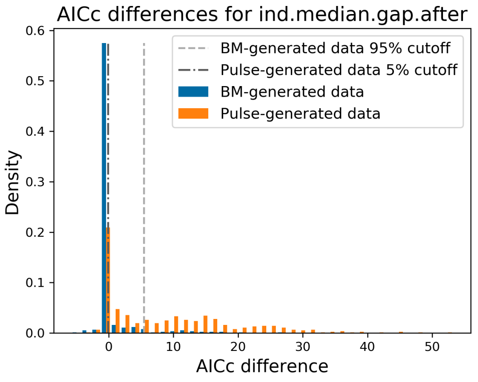
AICc differences for ind.median.gap.after
BM-generated data 95% cutoff
Pulse-generated data 5% cutoff
BM-generated data
Pulse-generated data
Density
AICc difference

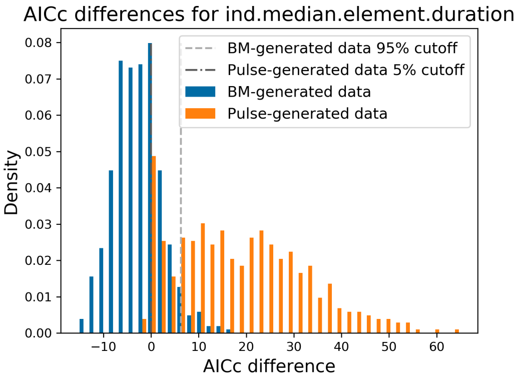
AICc differences for ind.median.element.duration
BM-generated data 95% cutoff
Pulse-generated data 5% cutoff
BM-generated data
Pulse-generated data
Density
AICc difference

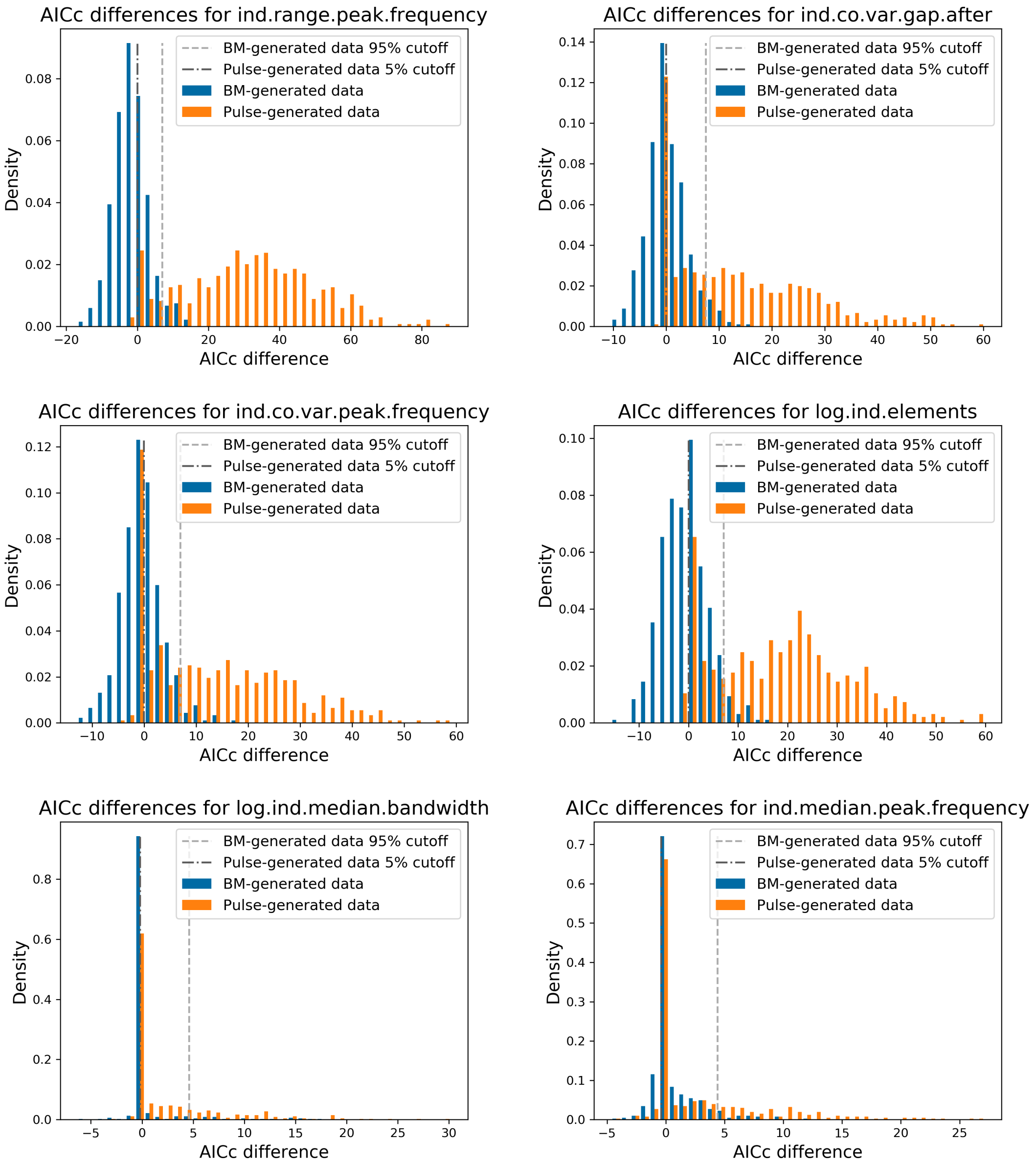


Figure S3: Empirical distribution of $\Delta A$ scores for pulse- and BM-generated artificial data, over parameter spaces relevant to each trait under consideration.

| Dataset | BM 5% | BM 95% | Pulse 5% | Pulse 95% | $\Delta A$ |
|---|---|---|---|---|---|
| CV frequency bandwidth | -5.611 | 6.943 | -0.160 | 23.757 | -1.173 |
| CV frequency change | -2.552 | 6.464 | -0.403 | 10.245 | 0.766 |
| max peak frequency (Hz) | -1.546 | 4.828 | -0.353 | 12.045 | 3.318 |
| log duration | -1.177 | 4.906 | -0.490 | 9.693 | 5.637 |
| min peak frequency (Hz) | -2.727 | 6.358 | -0.360 | 17.312 | 2.753 |
| log median frequency change | -0.465 | 5.456 | -0.092 | 28.747 | -0.001 |
| median pause duration | -10.486 | 6.264 | 0.000 | 43.496 | 5.610 |
| median element duration | -8.984 | 6.996 | 0.000 | 60.802 | 5.147 |
| range peak frequency | -5.474 | 7.483 | -0.033 | 40.605 | 11.610 |
| CV pause duration | -6.489 | 6.977 | -0.032 | 38.480 | 0.793 |
| CV peak frequency | -7.502 | 7.115 | -0.006 | 41.603 | 13.764 |
| number of elements | -0.215 | 4.601 | -0.195 | 13.328 | 3.630 |
| log median bandwidth | -1.229 | 4.405 | -0.398 | 14.654 | 1.784 |
| median peak frequency | -1.460 | 6.269 | -0.260 | 14.798 | 11.629 |

Table S2: $\Delta A$ 5th and 95th percentiles for data generated by the BM model (second and third column, respectively), and by the pulse model (fourth and fifth columns). Traits for which $\Delta A$ is greater than the 95th percentile of the empirical distribution of $\Delta A$ scores for BM-generated data are highlighted in dark gray. Traits which do not satisfy this constraint but are near the threshold, as suggested by the corresponding histograms in Figure S3, are highlighted in light gray.

### 4 Traits with pulses

#### 4.1 Pulse regimes and associated likelihoods

In addition to biological and statistical arguments, the plausibility of the minimal AICc pulse regimes can be observed in the correspondence between the ordered mean trait values and the different pulse regimes. In Figure S5, distinct clusters of values are shown placed in separate regimes. For these optimized pulse regimes, the optimality of the parameters may be verified from the likelihood surfaces (as functions of $\sigma_p^2$ and $\sigma_D^2$) given in Figure S6. Since these likelihood functions are conditioned on $\mu$ being optimal, we show dependence of $\mu$ on $\sigma_p^2$ and $\sigma_D^2$ separately in Figure S7.

#### 4.2 Pulse localization and weighing

In the case that the pulse model is favored, we quantify the support of a pulse on the $i$th branch by summing the AICc weights of the pulse configurations that have a pulse on that branch, as described in [2]. Since configurations with high support typically have similar parameter values, the corresponding constant corrections should be similar, and should therefore (approximately) cancel out in the weight computation. Pulse supports are given in the annotated phylogenetic trees in Figure S8, which were generated using the ETE Toolkit [6].

### 5 Sensitivity to tree topology

To characterize the uncertainty in model selection due to phylogenetic uncertainty, we built ten bootstrap population trees, and found the max likelihood parameters for the BM and pulse models. For each combination of model, tree and trait, we simulated 200 datasets (as above) with 1 pulse, and found the appropriate $\Delta A$ cutoff values using the 5th and 95th $\Delta A$ percentiles of the pulse- and BM-generated data. We then counted how many times each model was chosen for a given trait, using a different $\Delta A$ cutoff for each tree and trait combination.

For three of the four traits with strong evidence for pulsed evolution (range peak frequency, CV peak frequency, median peak frequency), the pulse model was selected under every tree. The fourth trait with strong evidence for pulsed evolution (log duration) was chosen under seven of the bootstrapped trees. Similarly, for two of the traits for which we found suggestive evidence for pulsed evolution, the pulse model was

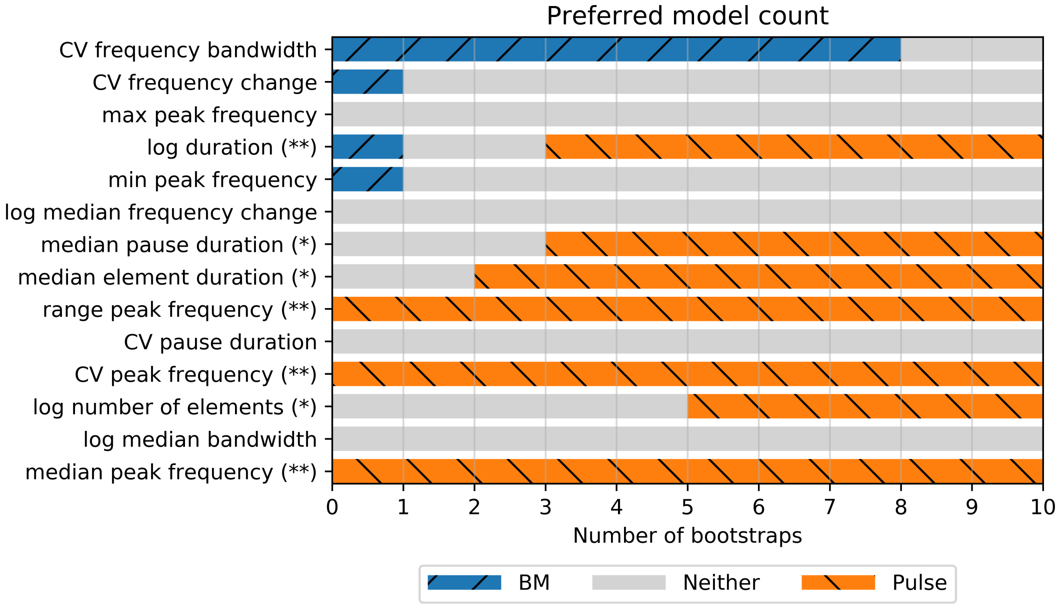


Figure S4: For each pair consisting of a data file and a bootstrapped tree, we fit both BM and pulse models, and estimate the cutoff $\Delta A$ values for model selection via 400 simulations. We then count the total number of trees for which each model is chosen. Traits for which there is strong support for the pulse model in Table S2 are denoted by two asterisks, and traits with suggestive support are denoted by one asterik. Pulse model selection appears consistent under tree perturbation.

chosen 7 and 8 times out of 10. For the trait of number of elements, the pulse model was selected for half of the trees, and no model was selected for the other half of the trees. These results in quantified in Figure S4.

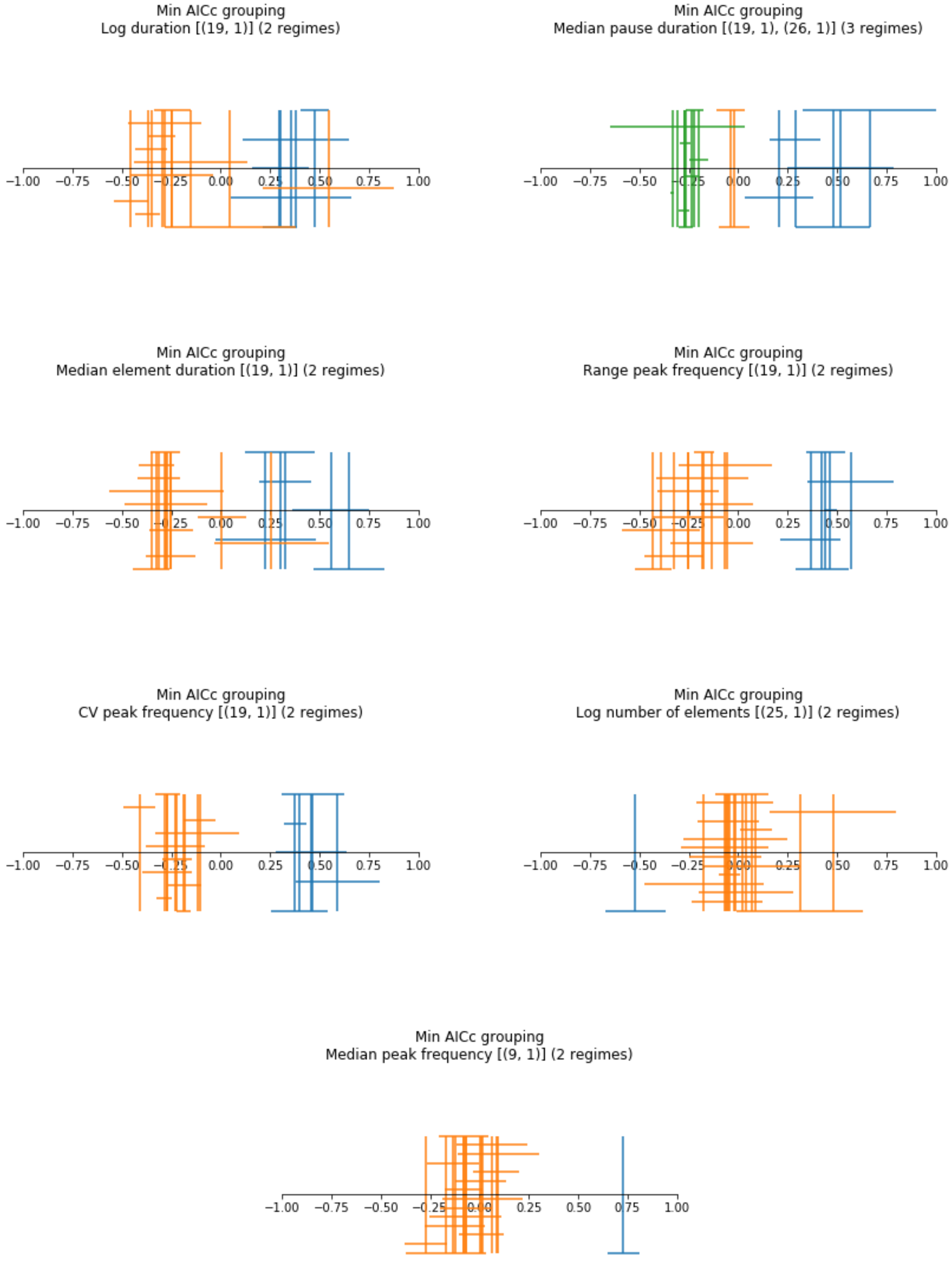


Figure S5: For each trait, each color group corresponds to different pulse regimes under the minimal AICc configuration. Only traits for which the pulse model is the preferred model are shown. Each vertical bar corresponds to a single population's mean value, and the horizontal bar centered on it denotes the standard error of each measurement.

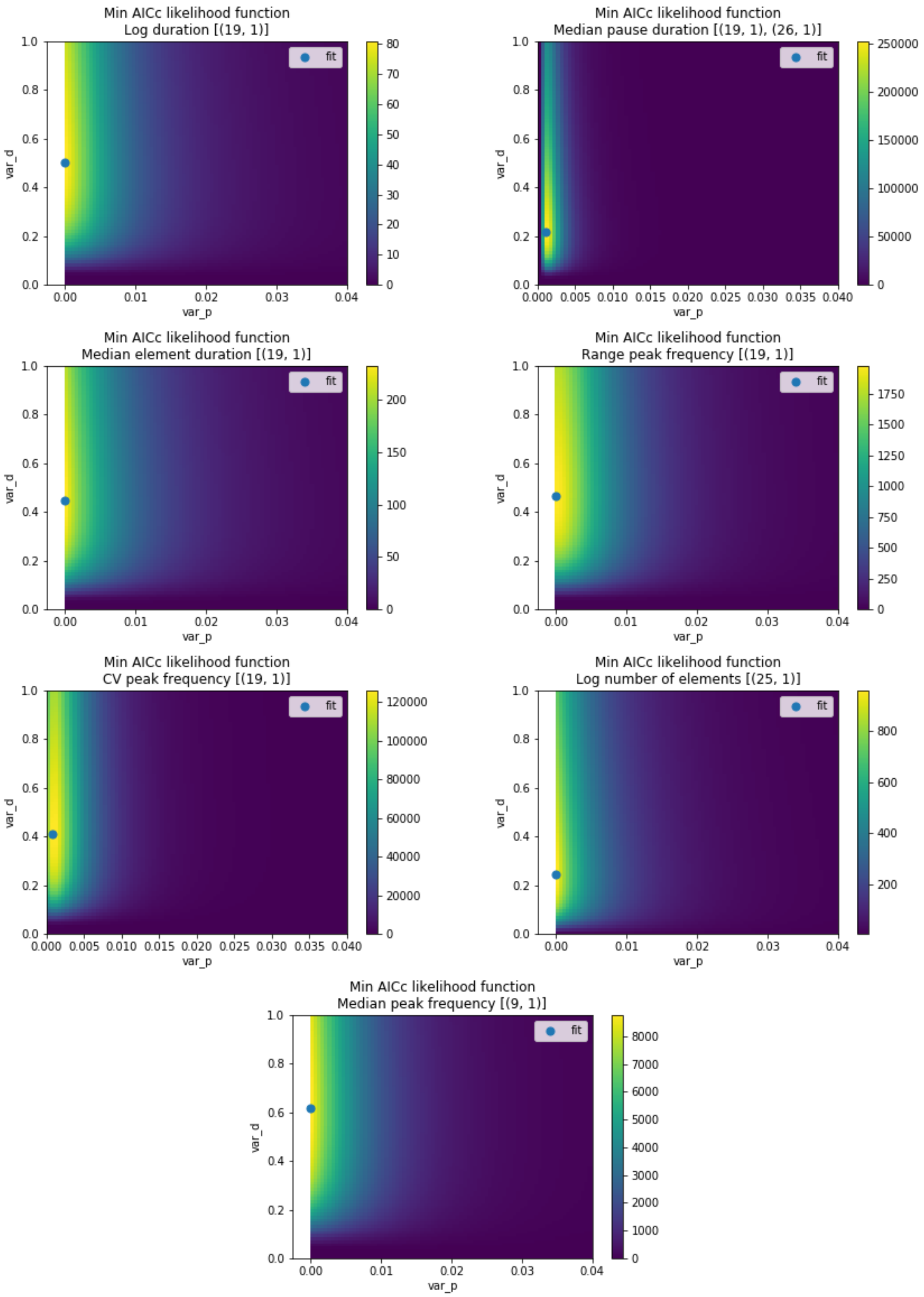


Figure S6: For each trait, we show the pulse model likelihood as a function of $\sigma_p^2$ and $\sigma_d^2$ under pulse configuration with lowest AICc. The trait mean $\mu$ is always taken as the optimal one for the given pair of variances. These graphs are therefore not simple cross-sections of the likelihood surface. The dependence of the mean on the variance parameters is shown in Figure S7. Only traits for which the pulse model has substantial evidence are shown.

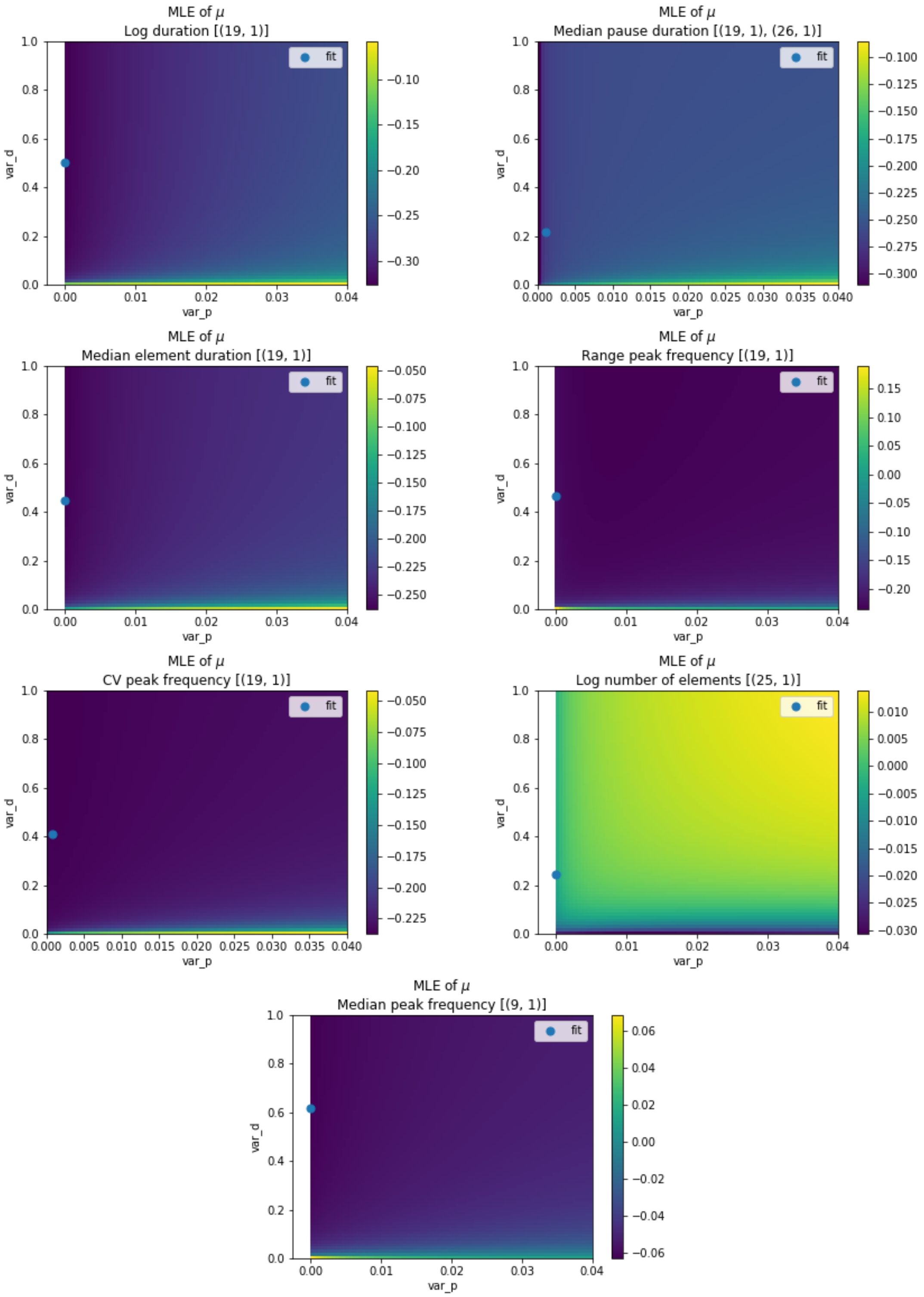


Figure S7: For each trait, we graph the ancestral mean as a function of $\sigma_p^2$ and $\sigma_d^2$ under minimal AICc pulse configuration. These are the values of $\mu$ used in Figure S6. Only traits for which the pulse model has substantial evidence are shown.

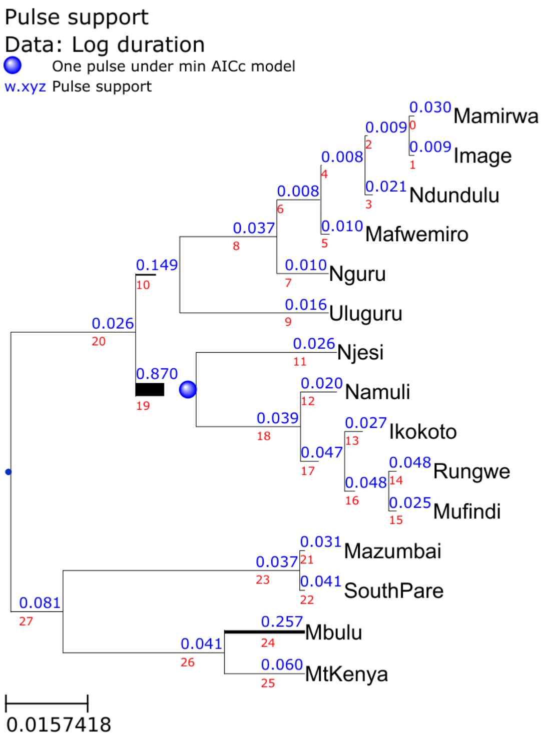
Pulse support
Data: Log duration
One pulse under min AICc model
w.xyz Pulse support
0.030 Mamirwa
0.009 Image
0.009
0.021 Ndundulu
0.008
0.010 Mafwemiro
0.008
0.010 Nguru
0.037
0.016 Uluguru
0.149
0.026
0.026 Njesi
0.870
0.020 Namuli
0.039
0.027 Ikokoto
0.047
0.048 Rungwe
0.048
0.025 Mufindi
0.031 Mazumbai
0.037
0.041 SouthPare
0.081
0.257 Mbulu
0.041
0.060 MtKenya
0.0157418

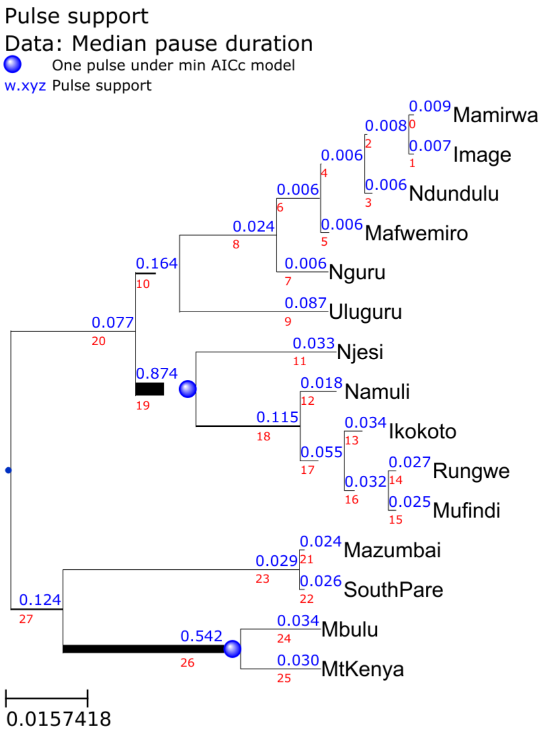
Pulse support
Data: Median pause duration
One pulse under min AICc model
w.xyz Pulse support
0.009 Mamirwa
0.008
0.007 Image
0.006
0.006 Ndundulu
0.006
0.006 Mafwemiro
0.024
0.006 Nguru
0.164
0.087 Uluguru
0.077
0.033 Njesi
0.874
0.018 Namuli
0.115
0.034 Ikokoto
0.055
0.027 Rungwe
0.032
0.025 Mufindi
0.024 Mazumbai
0.029
0.026 SouthPare
0.124
0.034 Mbulu
0.542
0.030 MtKenya
0.0157418

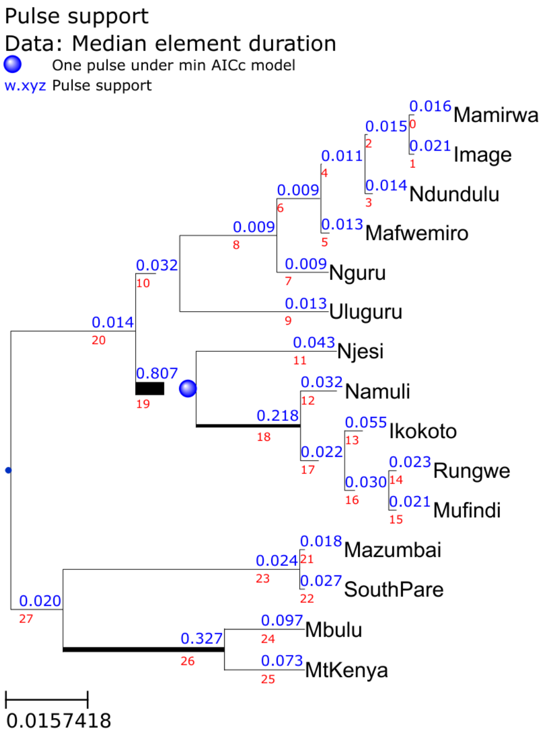
Pulse support
Data: Median element duration
One pulse under min AICc model
w.xyz Pulse support
0.016 Mamirwa
0.015
0.021 Image
0.011
0.014 Ndundulu
0.009
0.013 Mafwemiro
0.009
0.009 Nguru
0.032
0.013 Uluguru
0.014
0.043 Njesi
0.807
0.032 Namuli
0.218
0.055 Ikokoto
0.022
0.023 Rungwe
0.030
0.021 Mufindi
0.018 Mazumbai
0.024
0.027 SouthPare
0.020
0.097 Mbulu
0.327
0.073 MtKenya
0.0157418

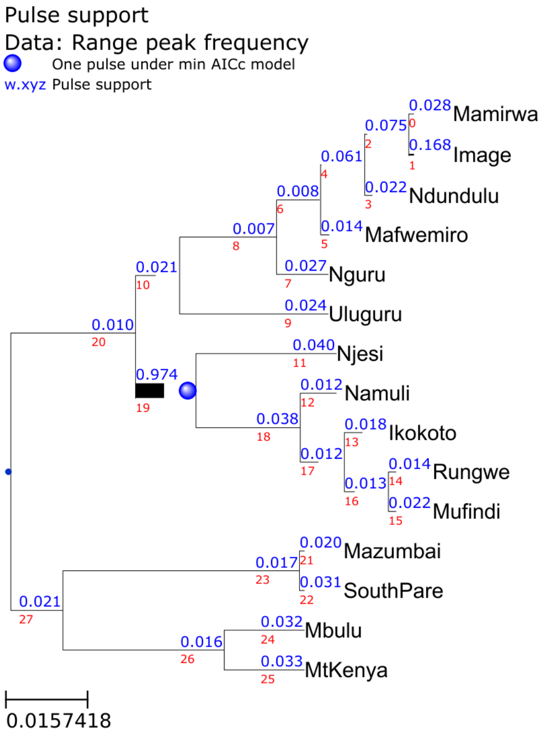
Pulse support
Data: Range peak frequency
One pulse under min AICc model
w.xyz Pulse support
0.028 Mamirwa
0.075
0.168 Image
0.061
0.022 Ndundulu
0.008
0.014 Mafwemiro
0.007
0.027 Nguru
0.021
0.024 Uluguru
0.010
0.040 Njesi
0.974
0.012 Namuli
0.038
0.018 Ikokoto
0.012
0.014 Rungwe
0.013
0.022 Mufindi
0.020 Mazumbai
0.017
0.031 SouthPare
0.021
0.032 Mbulu
0.016
0.033 MtKenya
0.0157418

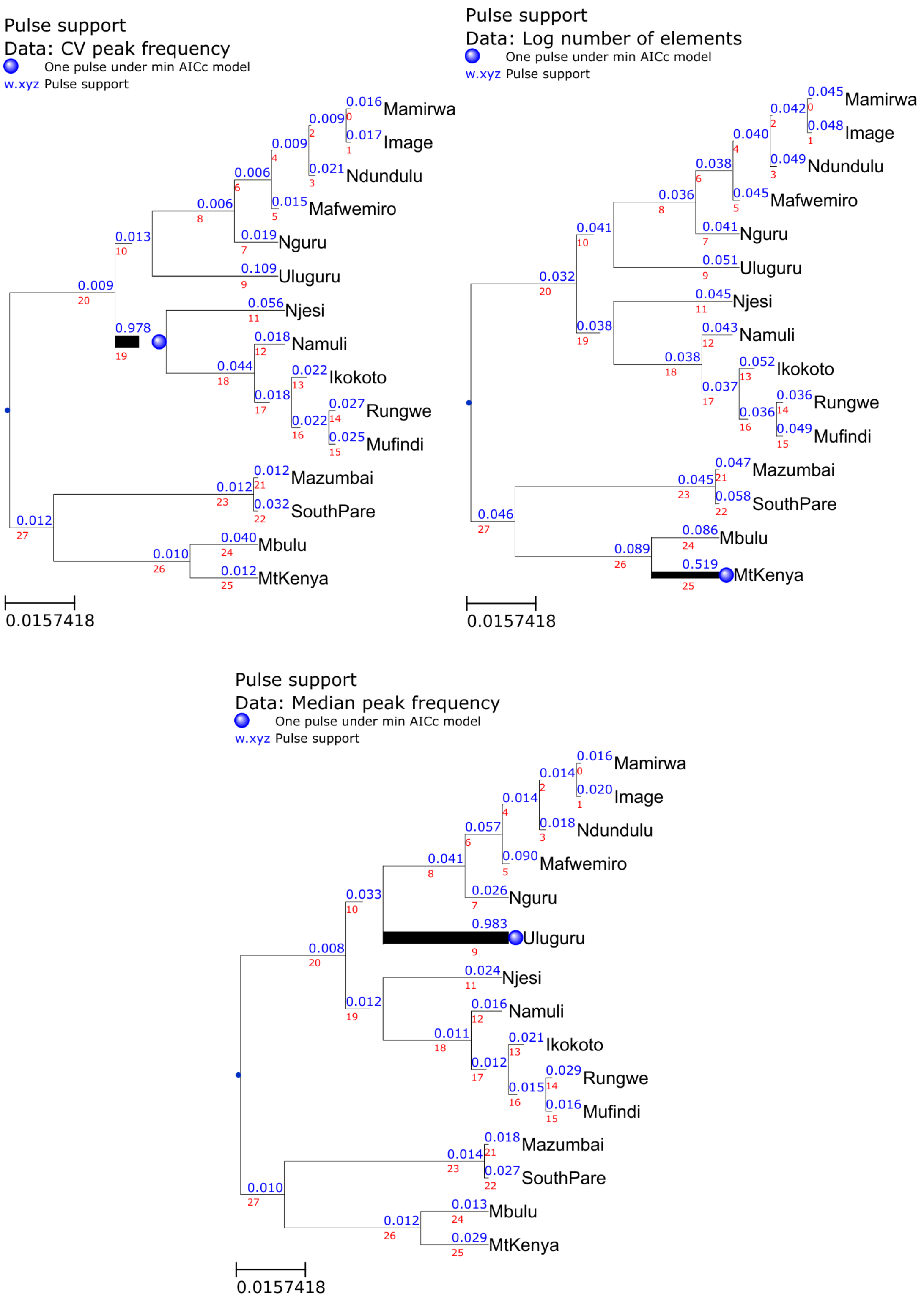


Figure S8: Pulse support on every branch. Only traits for which the pulse model has substantial evidence are shown.

### References


[1] Hirotugu Akaike. A new look at the statistical model identification. *IEEE Transactions on Automatic Control*, 19:716–723, 1974.

[2] Steven T Buckland, Kenneth P Burnham, and Nicole H Augustin. Model selection: an integral part of inference. *Biometrics*, pages 603–618, 1997.

[3] Kenneth P Burnham and David R Anderson. Multimodel inference: understanding AIC and BIC in model selection. *Sociological methods & research*, 33(2):261–304, 2004.

[4] Joseph Felsenstein. Maximum-likelihood estimation of evolutionary trees from continuous characters. *American journal of human genetics*, 25(5):471, 1973.

[5] Joseph Felsenstein. Phylogenies and the comparative method. *The American Naturalist*, 125(1):1–15, 1985.

[6] Jaime Huerta-Cepas, François Serra, and Peer Bork. ETE 3: reconstruction, analysis, and visualization of phylogenomic data. *Molecular biology and evolution*, 33(6):1635–1638, 2016.

[7] Clifford M Hurvich and Chih-Ling Tsai. Regression and time series model selection in small samples. *Biometrika*, 76(2):297–307, 1989.

[8] Clifford M Hurvich and Chih-Ling Tsai. Model selection for extended quasi-likelihood models in small samples. *Biometrics*, pages 1077–1084, 1995.

[9] Michael J Landis and Joshua G Schraiber. Pulsed evolution shaped modern vertebrate body sizes. *Proceedings of the National Academy of Sciences*, 114(50):13224–13229, 2017.

[10] Nariaki Sugiura. Further analysts of the data by Akaike's information criterion and the finite corrections: Further analysts of the data by Akaike's. *Communications in Statistics-Theory and Methods*, 7(1):13–26, 1978.
